# Enhanced nonenzymatic RNA copying with *in-situ* activation of short oligonucleotides

**DOI:** 10.1101/2023.04.12.536642

**Authors:** Dian Ding, Stephanie J. Zhang, Jack Szostak

## Abstract

The nonenzymatic copying of RNA is thought to have been necessary for the transition between prebiotic chemistry and ribozyme-catalyzed RNA replication in the RNA World. We have previously shown that a potentially prebiotic nucleotide activation pathway based on phospho-Passerini chemistry can lead to the efficient synthesis of 2-aminoimidazole activated mononucleotides when carried out under freeze-thaw cycling conditions. Such activated nucleotides react with each other to form 5′-5′ 2-aminoimidazolium bridged dinucleotides, enabling template-directed primer extension to occur within the same reaction mixture. However, mononucleotides linked to oligonucleotides by a 5′-5′ 2-aminoimidazolium bridge are superior substrates for nonenzymatic primer extension, due to their higher intrinsic reactivity and their higher template affinity. Here we show that eutectic phase phospho-Passerini chemistry efficiently activates short oligonucleotides and promotes the formation of monomer-bridged-oligonucleotide species during freeze-thaw cycles. We then demonstrate that *in-situ* generated monomer-bridged-oligonucleotides lead to efficient nonenzymatic template copying in the same reaction mixture. Our findings pave the way for future research into the activation of complex mixtures of mono- and oligonucleotides for the enhanced copying and potentially the replication of arbitrary RNA sequences.

## INTRODUCTION

The origin of life on the early Earth is likely to have involved a series of steps during which ribonucleotides were synthesized through abiotic chemical pathways, and subsequently assembled into short RNA oligonucleotides that encoded the genetic information of primordial life (1–6). A key step in the transition from prebiotic chemistry to the emergence of self-replicating RNAs and the advent of Darwinian evolution is thought to have been nonenzymatic RNA copying. For decades, template-directed nonenzymatic primer extension reactions have been modeled using intrinsically reactive mononucleotide phosphorimidazolides (Figure S1A) (7–9). Recent results suggest that 5′,5′-imidazolium-bridged-dinucleotides are the dominant reactive intermediates in facilitating chemical RNA copying in the presence of activated mononucleotides, due to their higher reactivity and stronger binding to the template by two Watson–Crick base pairs (Figure S1B) (10–13). Crystallographic studies have shown that the template-bound bridged dinucleotide intermediate is pre-organized so as to favor the primer extension reaction, such that the 3′-OH of the primer is properly oriented and in close proximity to the adjacent phosphate of the bridged dinucleotide (14, 15). While activated mononucleotides can contribute to template copying if present in high concentrations, the copying fidelity is extremely poor (16). Despite these advances, the chemical copying of RNA templates with all four canonical nucleotides remains inefficient. This inefficiency stems in part from the large differences in affinities and reactivities of the 10 different 2-aminoimidazole(2AI)-bridged dinucleotides, which results in low yields and sequence-dependent biases in template copying (13).

To overcome this problem, monomer-bridged-oligonucleotide intermediates have been proposed to facilitate fast and unbiased copying of templates containing all four bases (13, 17). Compared to activated mononucleotides and bridged dinucleotides, monomer-bridged-oligonucleotides can bind to the primer-template complex with significantly higher affinity and lead to a 10-fold higher rate of primer extension at template saturation (Figure S2A) (13). Thus, short oligonucleotides of only 2 to 4 nucleotides in length can act as sequence-specific catalysts for efficient template copying. Such short oligonucleotides may be generated prebiotically via untemplated (18–20) or template-directed oligomerization (21–23). In addition to primer extension, template copying can also be driven by the ligation of short oligomeric substrates, which is slower but requires fewer reaction steps (24–25). The nonenzymatic templated assembly of functional ribozymes has been shown to be possible using splinted ligation (25–27). Therefore, it is important to explore whether monomer-bridged-oligonucleotide substrates might also enhance nonenzymatic RNA ligation for template copying (Figure S2B).

Given the enhanced reactivity of monomer-bridged-oligonucleotides and the likely prebiotic availability of short oligonucleotides, we have investigated the *in-situ* activation of short oligonucleotides to see if they can be utilized as prebiotically available catalysts. Sutherland and coworkers have previously reported a nucleotide activation chemistry utilizing methyl isocyanide (MeNC), a compound that can be synthesized in a prebiotic ferrocyanide- and nitroprusside-containing environment after ultraviolet (UV) irradiation (28). By modifying this MeNC-mediated phospho-Passerini activation chemistry, our lab has recently identified a potentially prebiotic pathway for the activation of RNA mononucleotides and the formation of bridged dinucleotides under conditions compatible with nonenzymatic template copying (29). This reaction is conducted in the eutectic phase of a partially frozen reaction mixture to increase the effective concentrations of reactants, and thus promote bridged dinucleotide formation at a stoichiometric 2AI concentration. This pathway allowed us to demonstrate a prebiotically plausible scenario in which the hitherto separated stages of nucleotide activation, bridged dinucleotide formation, and nonenzymatic RNA copying occurred concurrently under mutually compatible conditions. Considering the expected heterogeneity of prebiotic mixtures of RNA mono- and oligonucleotides, we have now investigated the activation of short oligonucleotides and explored conditions that allow the subsequent formation of monomer-bridged-oligonucleotides for enhanced nonenzymatic RNA copying.

Here we show that monomer-bridged-oligonucleotides enhance both nonenzymatic primer extension and ligation. We then demonstrate that even low concentrations of activated oligonucleotides can generate sufficient monomer-bridged-oligonucleotide intermediates to drive efficient template copying. Following the demonstration of enhanced RNA copying with pre-activated oligonucleotides, we employ the MeNC-mediated chemistry to activate oligonucleotides and drive the formation of monomer-bridged-oligonucleotides. Finally, we demonstrate efficient nonenzymatic template copying starting with unactivated mono- and oligonucleotides under conditions of *in-situ* activation.

## MATERIAL AND METHODS

### General information

Unless otherwise noted, all chemicals were purchased at the highest available purity from Sigma-Aldrich (St. Louis, MO). All ribonucleoside 5′-monophosphates were obtained as free acids from MP Biotechnology (Solon, OH) or Sigma-Aldrich (St. Louis, MO). 2-Aminoimidazole hydrochloride was obtained from Combi-Blocks (San Diego, CA). Phosphoramidites and reagents for solid-phase RNA synthesis were purchased from ChemGenes (Wilmington, MA) and Glen Research (Sterling, MA).

Deuterated solvents were purchased from Cambridge Isotope Laboratories (Tewksbury, MA). The synthesis and storage of methyl isocyanide were carried out as previously described (30). To prevent the acidic hydrolysis of MeNC, which forms methylamine and formic acid, MeNC should be stored at a pH above 8. The absolute concentrations of stock solutions were determined by comparing the integrals of ^1^H peaks of interest to the calibrant, adenosine-5′-monophosphate, using NMR spectroscopy. The exact concentrations of the nucleoside-5′-monophosphates were determined by the analysis of serial dilutions on a spectrophotometer. When pipetting small volumes of the relatively volatile MeNC, it is common to experience loss of the MeNC inside the pipette tip, likely due to rapid evaporation. To ensure that the correct amount of MeNC is transferred to the reaction mixture, we pre-wet the pipette tips (Sorenson, BioScience, Inc.) by pipetting MeNC up and down at least three times, followed by immediate transfer of the pipetted MeNC solution into the reaction mixture and vortexing to ensure complete mixing.

Reverse-phase flash chromatography was performed using prepacked RediSep Rf Gold C18Aq 50 g columns from Teledyne Isco (Lincoln, NE). Preparatory-scale high-performance liquid chromatography (HPLC) was carried out on an Agilent 1290 HPLC system equipped with an Agilent ZORBAX Eclipse-XDB C18 column (21.2 × 250 mm, 7 µm particle size) for reversed-phase chromatography. Analytical HPLC was carried out on an Agilent 1100 series with an Agilent Eclipse Plus C18 column (4.6 × 250 mm, 5 µm particle size). ^31^P NMR spectra were acquired at 25 °C on a Varian Oxford AS-400 NMR spectrometer (162 MHz for ^31^P).

### Oligonucleotide Synthesis

Primers and templates were purchased from Integrated DNA Technologies. The dinucleotide, trinucleotides, and tetranucleotide (pCG, pCGC, pACG, and pCGCA) used for activation chemistry were prepared in-house by solid-phase synthesis on a MerMade 6 DNA/RNA synthesizer (Bioautomation, Plano, TX). Authentic samples of the three riboadenosine dinucleotides 5′-5′-pyrophosphate di-adenosine (AppA), di-adenosine 5′-monophosphate with a 2′-5′ phosphodiester linkage (pA-2′-pA), and di-adenosine 5′-monophosphate with a 3′-5′ phosphodiester linkage (pA-3′-pA) were prepared by solid-phase synthesis on an Expedite 8909 DNA/RNA synthesizer. The synthesized oligonucleotides were then deprotected and purified by reverse-phase chromatography with a 50 g C18Aq column using a gradient elution of 0–10% acetonitrile over 10 column volumes (CVs) in 20 mM triethylammonium bicarbonate (TEAB) at pH 7.5.

### Preparation of activated mononucleotides, bridged dinucleotides, activated oligonucleotides, and mononucleotide-bridged-oligonucleotides

All 2-aminoimidazole-activated RNAs (*pN_n_ and Np*pN_n,_ n = 1, 2, 3, 4) were synthesized as previously described (13). The activated mononucleotide *pA was purified by reverse phase chromatography with a 50 g C18Aq column over 10 CVs of 0–10% acetonitrile in 2 mM TEAB (pH 8). The fractions containing *pA were adjusted to pH 9.5–10 with NaOH, aliquoted, and lyophilized. Apart from using 20 mM TEAB (pH 7.5), the activated oligonucleotides (*pCG, *pCGC, and *pCGCA) were purified in the same manner. The bridged dinucleotide Ap*pC was purified by reverse-phase HPLC on a C18 column with a gradient of 2–8 % acetonitrile over 20 min in 2 mM TEAB (pH 8) at a flow rate of 15 mL/min. Fractions containing Ap*pC were adjusted to pH 8 with HCl, aliquoted, and lyophilized. The monomer-bridged oligonucleotide intermediates Ap*pCG, Ap*pCGC, and Ap*pCGCA were purified by reverse-phase HPLC on a C18 column with a gradient of 2–10% acetonitrile over 27 min in 20 mM TEAB (pH 7.5) at a flow rate of 15 mL/min. Fractions containing the desired products were adjusted to pH 8 with HCl, aliquoted, and lyophilized.

### Nonenzymatic primer extension and ligation with pre-synthesized activated RNAs

FAM-labeled primer /FAM/AGUGAGUAACUC was used in the nonenzymatic primer extension and ligation reactions with pre-synthesized activated RNAs unless otherwise noted. The primer extension reactions used the template 5′-OH-<u>UGCGU</u>GAGUUACUCACUAAA. Ligation reactions used the template 5′-OH-<u>UGCG</u>GAGUUACUCACUAAA. Underlined sequences indicate the template region available for substrate binding.

*Kinetics of primer extension with a single activated RNA substrate (*pN*_*n*_ *or Np*pN*_*n*_). The primer-template complex was first prepared in an annealing solution containing 7.5 μM primer, 12.5 μM template, 50 mM Tris-HCl (pH 8), 50 mM NaCl, and 1 mM ethylenediaminetetraacetic acid (EDTA) by heating at 85 °C for 30 s and slowly cooling down to 25 °C at a rate of 0.1 °C/s. The annealed product was then diluted into the final reaction mixture to a final concentration of 1.5 μM primer, 2.5 μM template, 200 mM Tris-HCl (pH 8), 100 mM MgCl_2_, and the indicated concentration of the pre-activated substrate. The stock solutions of pre-activated species were prepared freshly at various concentrations immediately before being added to the final reaction mixture to initiate the template copying reaction at room temperature. At each time point, 0.5 μL of the reaction mixture was added to 25 μL of a quenching buffer containing 25 mM EDTA, 1X TBE, and 4 µM DNA strands complementary to the template in formamide. Template copying products were resolved by 20% (19:1) denaturing PAGE, scanned with a Typhoon 9410 scanner, and quantified using the ImageQuant TL software. Reactions containing a blocker oligonucleotide were performed as described above, except using the sequences listed in Figure S4D, with final concentrations of 1.5 μM primer, 2.5 μM template, and 3.5 μM blocker.

*Primer extension with a mixture of activated mono- and oligonucleotides (*pA and *pN*_*n*_). The annealing buffer was prepared as described above with 7.5 μM primer and 12.5 μM template. The annealing buffer containing the primer-template complex was mixed with MgCl_2_ and Tris-HCl (pH 8) at the bottom of the reaction tube, while the freshly prepared stock solutions of activated mononucleotide (*pA) and activated oligonucleotide (*pCG, *pCGC, or *pCGCA) were added separately to the lid or wall of the tube. The reaction tube was immediately centrifuged and vortexed to mix all the materials to initiate the reaction with final concentrations of 1.5 μM primer, 2.5 μM template, 200 mM Tris-HCl (pH 8), 100 mM MgCl_2_, 5 mM *A, and 5/0.5/0.05 mM activated oligonucleotide. Samples were collected at each time point and analyzed as described above.

*Primer extension with a pre-incubated mixture of activated mono- and oligonucleotides (*pA and *pCGCA)*. The annealed primer-template complex was prepared as described above and lyophilized to dryness. A mixture of 5 mM *A, 0.5 mM *CGCA, 200 mM Tris-HCl (pH 8), and 100 mM MgCl_2_ was incubated for 1 h at room temperature before being added to the lyophilized primer-template complex, to give final concentrations of 1.5 μM primer and 2.5 μM template. Samples were collected at each time point and analyzed as described above.

### HPLC analysis of spontaneous bridge-formation between activated mono- and oligonucleotides

The activated tetramer *pCGCA was used for the kinetic analysis of bridged species due to its strong UV absorbance, which allowed for effective analysis by HPLC. In primer extension buffer (100 mM MgCl_2_ & 200 mM Tris-HCl (pH 8) or 30 mM MgCl_2_ & 50 mM Na^+^-Hepes (pH 8)), 5 mM *pA was allowed to react with 0.5 mM *pCGCA. After incubation at room temperature for the desired time, 0.5 M EDTA solution was added in 2.5-fold excess over the MgCl_2_ concentration, and the resultant mixture was kept on dry ice before injection or injected immediately into an Agilent ZORBAX Eclipse-X DB C18 column for HPLC analysis. The sample was separated using (A) aqueous 25 mM TEAB buffer (pH 8) and (B) acetonitrile, with a gradient of 3% to 10 % B over 25 min at a flow rate of 15 mL/min. Fractions were collected and analyzed by mass spectroscopy to confirm the identity of the collected species.

### HPLC analysis of MeNC-mediated activation and bridge-formation

#### Oligonucleotide activation at ambient temperature

400 mM MeNC, 400 mM 2-methylbutyraldehyde (2MBA), and 200 mM 2AI were added to a solution of 5 mM oligonucleotides (2-4 nucleotides in length) in 200 mM Na^+^-Hepes at pH 8 with 30 mM Mg^2+^. The reaction was allowed to sit for 6 hours, the optimal incubation time previously determined (30). The mixture was then either brought to 10 % (v/v) D_2_O for NMR spectroscopy, or separated from MeNC-mediated activation reagents using Sep-Pak^®^ C18 Cartridges for HPLC analysis. To perform the Sep-Pak^®^ cleanup, the stationary phase of the cartridge was first wetted with acetonitrile and 2 M triethylamine acetate. The reaction sample was then diluted in 1 mL of UltraPure DNase/RNase-free distilled water and slowly loaded onto the cartridge three times. The cartridge was then washed three times with 3 mL of 20 mM TEAB (pH 8) before the oligonucleotides were slowly eluted with 2 mL of 40% acetonitrile in 20 mM TEAB (pH 8). The concentration of the total oligonucleotides was determined by a NanoDrop Microvolume Spectrometer. Acetonitrile was removed by leaving the eluate under ambient temperature for 1 hour. The sample was then lyophilized, redissolved in UltraPure water, and analyzed on an Agilent 1100 series HPLC with an Agilent ZORBAX Eclipse-XDB C18 column. The sample was separated using (A) aqueous 25 mM TEAB buffer (pH 8) and (B) acetonitrile, with 2% B for 5 min, then 2% to 13% B over 30 minutes, unless otherwise noted. All peak fractions were flash-frozen and lyophilized before confirming their identity by liquid chromatography-mass spectrometry (LC-MS).

#### Eutectic phase activation of mononucleotides and oligonucleotides

A mixture of 5 mM mononucleotide and indicated concentrations oligonucleotides was allowed to react with stoichiometric 2AI ([2AI] = [pN]+[pN_n_]), 30 mM MgCl_2_, 50 mM Na^+^-Hepes (pH 8), 200 mM 2MBA, and 50 mM MeNC under eutectic ice phase conditions. Rapid cooling by liquid nitrogen was used to ensure complete freezing before the solution was stored at -15 to -13 °C. Every 24 h, the sample was thawed, 50 mM additional MeNC was added, and refrozen to the eutectic ice phase. After four days, the products were separated from MeNC-mediated activation reagents and analyzed by analytical HPLC as described above.

#### Nonenzymatic primer extension and ligation with *in-situ* activated RNAs

All reactions used the thiol-modified primer /5ThioMC6-D/AGUGAGUAACUC. The primer extension reactions used the template 5ʹ-OH-<u>UGCGU</u>GAGUUACUCACUAAA. Ligation reactions used the template 5ʹ-OH-<u>UGCG</u>GAGUUACUCACUAAA. Underlined sequences indicate the template region available for substrate binding.

All reactions contained 1 μM primer, 1.5 μM template, 50 mM Na^+^-Hepes (pH 8), and 30 mM MgCl2. Unactivated mononucleotides were supplied at 5 mM each, while unactivated pCGC was supplied at 0.5 mM, except as otherwise noted. 2AI was supplied at a stoichiometric concentration ([2AI] = [pN]+[pN_n_]), except for the cases containing only pA or pCGC, where excess 2AI was supplied at 5.5 mM. The primer and template were first mixed with Na^+^-Hepes (pH 8). Then stock solutions of unactivated mononucleotides and/or oligonucleotides, and 2AI were prepared separately, adjusted to pH 8 with NaOH or HCl, and added to the primer-template mixture, followed by the addition of MgCl_2_.

The reaction mixture was brought to 200 mM 2MBA and 50 mM MeNC before being frozen to the eutectic phase at approximately -15 to -13°C. The freezer temperature was monitored, and slight temperature fluctuations were observed occasionally. Rapid cooling by liquid nitrogen was used to ensure complete freezing before the sample was incubated at the ice-eutectic phase. Every 24 h, the mixture was thawed to room temperature for sample collection, an additional 50 mM MeNC was added, and then subjected to eutectic freezing for one more day. Thawing, sample collection, and refreezing took less than ten minutes. After four days of eutectic activation, the reaction mixture was brought to room temperature and allowed to react for another 24 h. The mixture was then incubated at ambient temperature for two more days with the addition of 100 mM 2MBA and 100 mM MeNC per day.

Every 24 h, 30 μL of the reaction mixture was collected and purified by ZYMO Oligo Clean & Concentrator spin column (ZYMO Research, Irvine, CA). The isolated material was then mixed with 30 μL 100 mM Na^+^-Hepes (pH 8), and any disulfides were reduced using a 10-fold molar excess of tris-(2-carboxyethyl) phosphine hydrochloride (TCEP) for 1.5 ∼ 2 h. Then 0.8 μL of 1 mM Alexa 488 C5 maleimide dye dissolved in anhydrous dimethyl sulfoxide was added to the reduced mixture, and the coupling reaction was allowed to proceed in the dark at ambient temperature for 1.5 ∼ 2 h. The labeled primer-template duplex was separated from free dyes using ZYMO DNA Clean & Concentrator spin columns and concentrated to a volume of 10 μL. 1 μL of the dye-labeled mixture was added to 3 μL of gel loading buffer containing 8 M Urea, 10X TBE, and 100 μM of an RNA complementary to the template. The sample was heated to 95 °C for 30 s and cooled to ambient temperature to ensure the separation of the template from the dye-labeled primer. Template copying products were resolved by 20 % (19:1) denaturing PAGE, scanned with a Typhoon 9410 scanner, and quantified using the ImageQuant TL software.

#### Rapid quenching of nonenzymatic primer extension with *in-situ* activated pA & pCGC

*Comparison between EDTA quenched and thawed reactions*. Parallel experiments were set up at the same time with identical contents in 40-μL scale: 5 mM pA, 0.5 mM pCGC, 5.5 mM 2AI, 1 μM primer (/5ThioMC6-D/AGUGAGUAACUC), 1.5 μM template (5ʹ-OH-<u>UGCGU</u>GAGUUACUCACUAAA), 50 mM Na^+^-Hepes (pH 8), and 30 mM MgCl_2_. The reaction mixtures were treated with 200 mM 2MBA and 50 mM MeNC and then immersed in liquid nitrogen to ensure complete freezing. Next, the samples were kept at - 15 to -13 °C for ice-eutectic phase reaction. After 24 hours, one of the frozen 40-μL samples was added to 10 μL of 0.5 M EDTA and vortexed to make sure all the thawed solution was immediately quenched by EDTA. Another frozen sample was allowed to thaw at ambient temperature. Then the two samples were purified by ZYMO Oligo Clean & Concentrator spin columns, reduced by TCEP and dye-labeled by Alexa 488 C5 maleimide dye, purified again, and resolved by 20% (19:1) denaturing PAGE gel following the same protocols as described earlier.

*Time course of the EDTA-quenched reaction*. Parallel experiments were set up as described above in a 40-μL scale for each. Each reaction was individually quenched by EDTA at the desired time point before thawing, and analyzed following the same procedure.

#### Nonenzymatic primer extension with eutectic-phase-activated RNAs

A mixture of 5 mM pA, 0.5 mM pCGC, 5.5 mM 2AI, 50 mM Na^+^-Hepes (pH 8), and 30 mM MgCl_2_ was brought to 200 mM 2MBA and 50 mM MeNC. Next, the mixture was immersed in liquid nitrogen to ensure complete freezing and then kept at -15 to -13°C for 24 h. The primer-template mixture was prepared and lyophilized into dry powder. The MeNC-activated mixture in the ice eutectic phase was thawed and immediately added to the primer-template powder to give a solution containing 1 μM primer (/5ThioMC6-D/AGUGAGUAACUC) and 1.5 μM template (5′-OH-<u>UGCGU</u>GAGUUACUCACUAAA). At each time point, 20 μL of the reaction mixture was added to 10 μL 0.5 M EDTA and purified by ZYMO Oligo Clean & Concentrator spin columns. The thiolated RNAs were reduced with TCEP and labeled by incubating with Alexa 488 C5 maleimide dye, purified again, and resolved by a 20% (19:1) denaturing PAGE gel as described above.

## RESULTS

### Kinetic analysis of RNA copying with pre-synthesized monomer-bridged-oligonucleotides

We started by asking whether monomer-bridged-oligonucleotides can significantly improve the efficiency of template-directed ligation as well as nonenzymatic primer extension. We have previously reported kinetic parameters for several nonenzymatic primer extension reactions using monomer-bridged-oligonucleotides (13). We have repeated these experiments, and compared the kinetics of primer extension with ligation under identical conditions (Figure 1A, S3). Our new measurements show almost identical Michaelis constants (K_M_) as proxies for binding affinity, compared to previously reported data. Thus, monomer-bridged-oligonucleotides bind to the primer-template complex much more strongly than activated mononucleotides or bridged dinucleotides. Furthermore, the maximum reaction rates (k_obs max_) for primer extension with *pA are increased by over 80-fold when *pA is bridged to a downstream mononucleotide pC, and by 500 to 700-fold when *pA is bridged to an oligonucleotide (Figure 1A). The enhanced reaction rate at saturation suggests a greater degree of pre-organization into an optimal configuration for reactions (15, 31).

**Figure 1.**
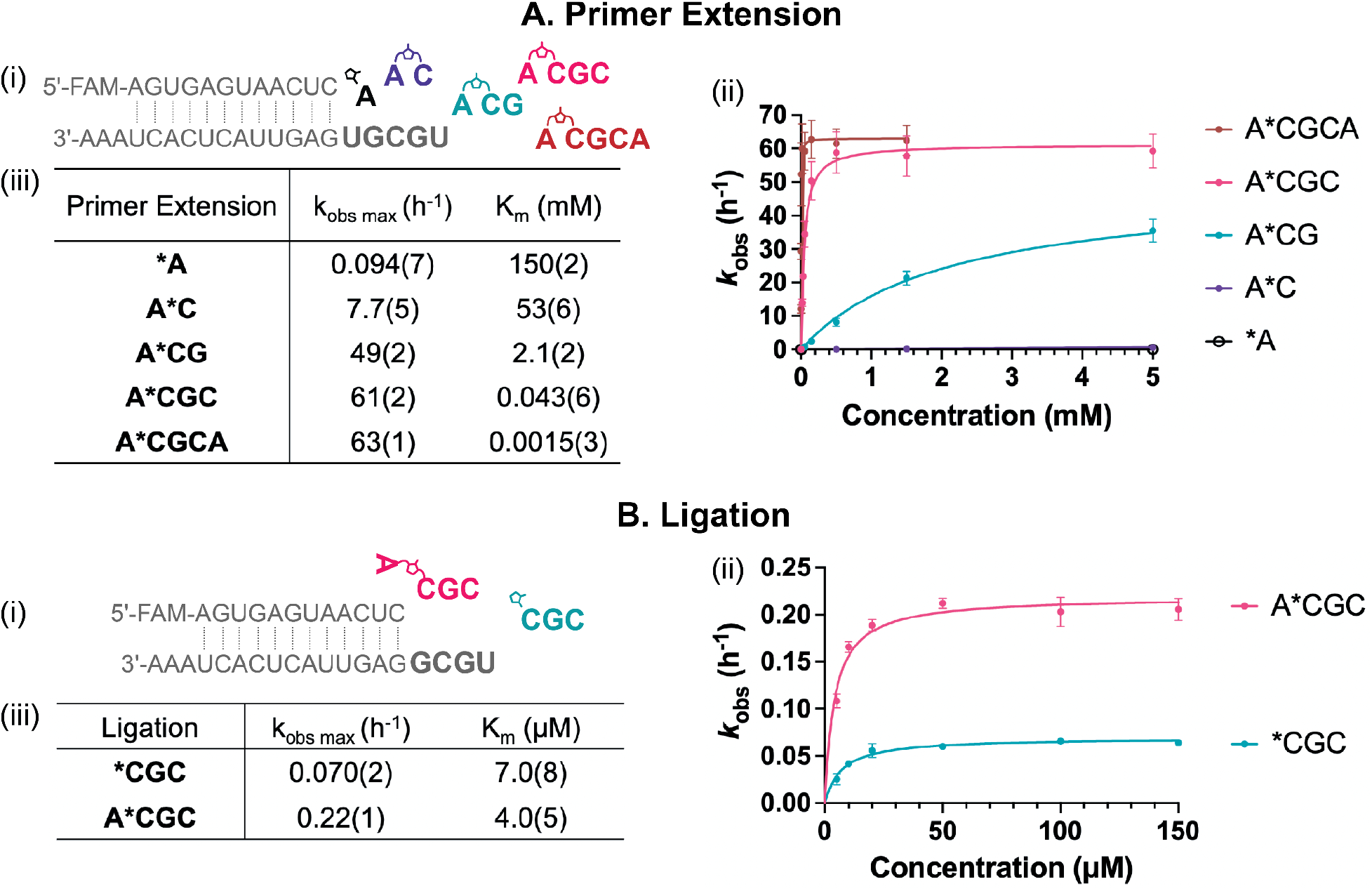
Kinetics of nonenzymatic primer extension and ligation with pure monomer-bridged-oligonucleotides. (A) Primer extension. (i) Schematic representation of nonenzymatic single nucleotide primer extension using a series of activated species. (ii) Michaelis-Menten analysis of primer extension with the illustrated activated species. The k_obs_ values for A*C and *A are too low to display properly in the figure; see Figure S3 for the full Michalis-Menten plot for each substrate. (iii) Kinetic parameters for +1 primer extension. Modified from Fig. 7 in ref. 14 with permission under a Creative Commons Attribution 4.0 International License. Copyright 2022 Ding et al.; Published by Oxford University Press on behalf of Nucleic Acids Research. (B) Ligation. (i) Schematic representation of nonenzymatic ligation with either an activated trinucleotide or a monomer-bridged-trinucleotide. (ii) Michaelis-Menten plots for nonenzymatic ligation. (iii) Kinetic parameters for ligation. All reactions were performed with 100 mM MgCl_2_ and 200 mM Tris-HCl, pH 8 at ambient temperature.

Given the enhanced nonenzymatic primer extension observed with the monomer-bridged-oligonucleotides, we next asked whether the imidazolium bridging moiety, a stronger electron withdrawing group than imidazole, would also accelerate ligation in the same conditions (100 mM MgCl_2_, 200 mM Tris-HCl, pH 8) when the bridged mononucleotide component is not bound to the template (Figure 1Bi, S2B). In this configuration, and under the same reaction conditions, we found that the maximum ligation rate of Ap*pCGC was more than three-fold higher than that of *pCGC (Figure 1B). To further explore the effect of an unbound imidazolium-bridged nucleotide, we employed a sandwiched primer-template-blocker system with a single-nucleotide binding site on the template, and let the primer extend with either *pN or Np*pN (Figure S4A). We found that an imidazolium-bridged dinucleotide in which only one nucleotide is bound to the template exhibits a more than a five-fold increase in the k_obs max_ of +1 primer extension reaction (100 mM MgCl_2_, 200 mM Tris-HCl, pH 8). This rate enhancement applies to all four canonical nucleotides as the leaving group (Figure S4). These observations suggest that the superior electrophilicity of the imidazolium bridging unit can enhance both primer extension and ligation. While decreasing the reaction pH to protonate the imidazole group of an activated oligonucleotide may also generate a positively charged leaving group just like the imidazolium bridging moiety, deprotonation of the primer 3ʹ-OH (as shown in Figure S1) would be more challenging at this lower pH, making the nucleophilic attack less favorable (32). By employing the imidazolium bridging moiety as the more electrophilic leaving group, we were able to achieve faster nonenzymatic ligation under the same conditions as used for primer extension, while using essentially the same reactive intermediates. Additionally, this approach could potentially alleviate the inhibition of primer extension caused by the binding of long RNAs downstream of a primer. By activating the 5′-end of the blocking oligonucleotide with an imidazolium-bridged nucleotide, this blocking strand can take part in faster nonenzymatic ligation.

### Chemical RNA copying with a mixture of activated mono- and oligonucleotides

The significantly improved RNA copying observed with monomer-bridged-oligonucleotide substrates prompted us to consider a more prebiotically plausible scenario in which activated mononucleotides were present together with activated oligonucleotides. To investigate this, we performed experiments with a fixed concentration of 5 mM of *pA, and lower concentrations of downstream oligonucleotides 2, 3 and 4 nucleotides in length. The *pA and activated oligonucleotides react with each other in solution to form the monomer-bridged-oligonucleotide substrates for primer extension. We find that as little as 50 μM activated trinucleotides or tetranucleotides is sufficient to enhance the yield of primer extension, whereas the activated dimer had to be present at a higher concentration (of 5 mM) to achieve a similar yield (Figure 2A-C). However, the primer extension yield as a function of time exhibits a lag characteristic of a two-step reaction, indicating a requirement for the initial formation of monomer-bridged-oligonucleotides before significant primer extension could occur. Since two sequential reactions, bridge formation and primer extension, happen in this scenario, it was difficult to quantitatively assess only the rate of the later primer extension reaction.

**Figure 2.**
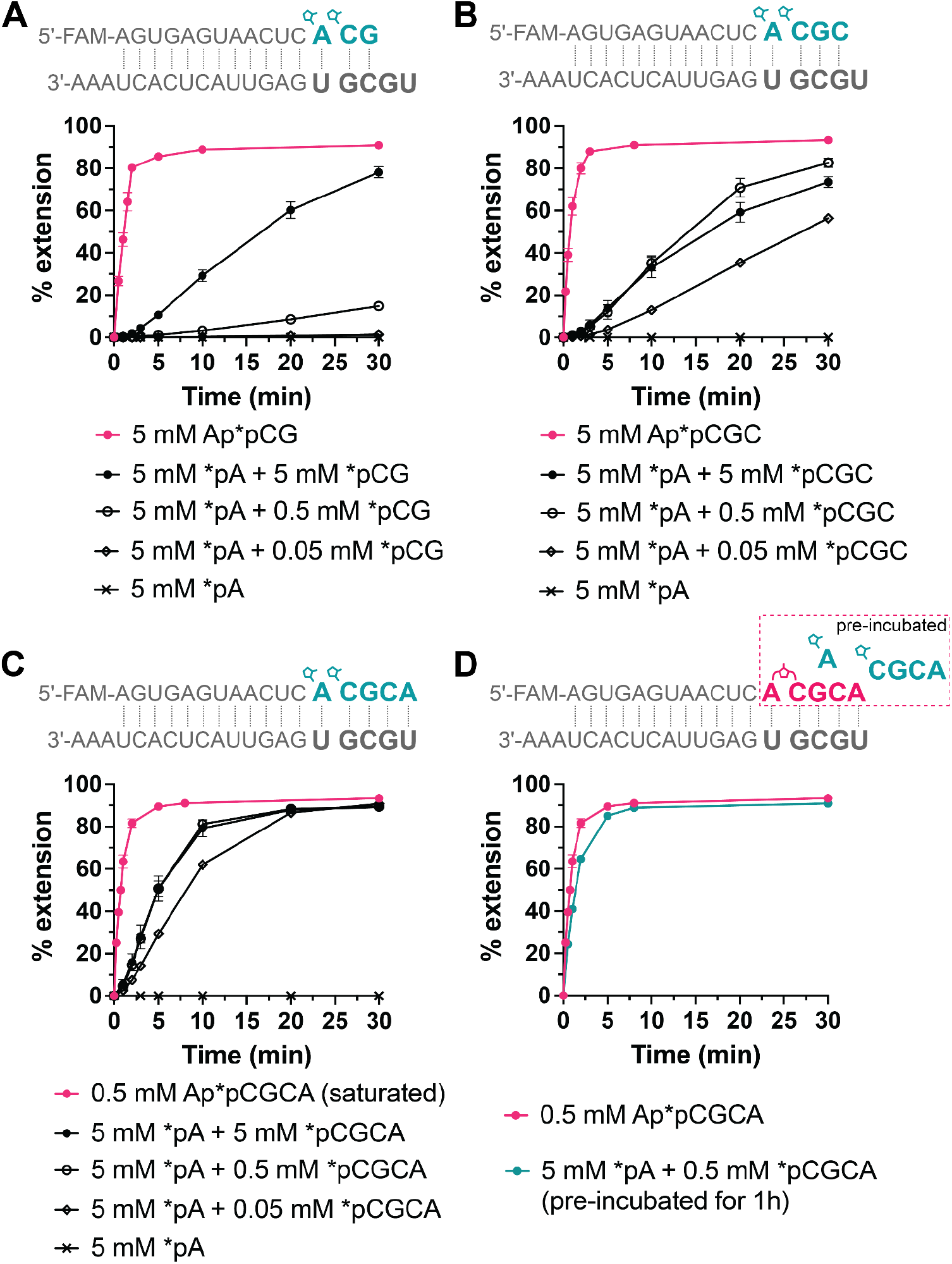
Comparison of primer extension with pre-synthesized monomer-bridged-oligonucleotide vs. mixtures of pre-activated mono- and oligonucleotides. (A) Comparison of primer extension with Ap*pCG and *pA + *pCG. (B) Comparison of primer extension with Ap*pCGC and *pA + *pCGC. (C) Comparison of primer extension with Ap*pCGCA and *pA + *pCGCA. (D) Comparison between Ap*pCGCA and a mixture of *pA + *pCGCA preincubated for 1 h. Negative controls with only 5 mM *pA are shown in A-C. No detectable extension with *pA was observed in the first hour. All reactions were performed with 100 mM MgCl_2_ and 200 mM Tris-HCl, pH 8.

In order to observe the primer extension reaction without the initial delay, we pre-incubated mixtures of activated mono- and oligonucleotides and measured the time required for spontaneous monomer-bridged-oligonucleotide formation in solution. A mixture of 5 mM *pA and 0.5 mM *pCGCA was allowed to react in a primer extension buffer (100 mM MgCl_2_ and 200 mM Tris-HCl, pH 8) at room temperature, and we measured the formation of Ap*pCGCA at each time point by HPLC. We found that Ap*pCGCA spontaneously formed from the mixture of *pA and *pCGCA, with its concentration peaking at 1 h (∼53 μM, 11% conversion from *pCGCA to Ap*pCGCA, Figure S5). After 1 h, the concentration of Ap*pCGCA started to decline, likely due to the hydrolysis. To examine primer extension, we pre-incubated a mixture of 5 mM *pA and 0.5 mM *pCGCA in a primer extension buffer for 1 h before adding that to the primer-template complex. We were pleased to find that primer extension proceeded with no lag, and at a rate very similar to that observed with pre-synthesized Ap*pCGCA (Figure 2D). Our results suggest that activating even a small fraction of the oligonucleotides in a complex mixture may be sufficient to enable efficient nonenzymatic template copying.

### MeNC-mediated activation and bridge-formation of monomers and oligonucleotides

The efficient nonenzymatic RNA copying facilitated by modest concentrations of activated oligonucleotides prompted us to investigate the activation of oligonucleotides under potentially prebiotic conditions. We started by demonstrating that short oligonucleotides can be activated using MeNC, 2MBA, and excess 2AI in the same buffer used for nonenzymatic RNA copying (50 mM Na^+^-Hepes pH 8 and 30 mM MgCl_2_). When the MgCl_2_ concentration was reduced to suppress hydrolysis and side product formation (30), we found that short oligonucleotides of various lengths can be activated with good efficiency (Figure S6). For example, approximately 89% of dimers (pCA) and 77% of trimers (pCCA) were activated to 2-aminoimidazolides under these conditions. In addition to the efficient activation of oligonucleotides, our data suggest that the effective concentrations of reagents required for activation are remarkably similar for mononucleotides and oligonucleotides up to tetranucleotides.

Given the efficient solution phase activation of oligonucleotides with excess 2AI, we next explored whether 2AI-bridged mono- and oligonucleotides could form with stoichiometric 2AI using MeNC-mediated chemistry. To do this, we employed eutectic ice phase conditions, which we have previously shown to facilitate the efficient activation of mononucleotides and the formation of bridged dinucleotides. We began by activating 5 mM pA and 2 mM pCG under the eutectic ice phase (−15 to -13 °C) with 7 mM 2AI. After four days of activation, more than 60% of the pA and pCG were converted into either activated or bridged species (i.e., bridged dinucleotides or monomer-bridged-dinucleotides) as determined by HPLC analysis (Figures 3B and 3Ci). Actual levels of activation and bridge formation in the reaction mixture may be higher because of hydrolysis during post-reaction cleanup. Side products, including the 3′-5′ linked dinucleotide, were minimal, as confirmed by the spike-in of their corresponding authentic standards (Figure S7). We then proceeded to activate longer oligonucleotides at decreasing concentrations because of their possible lower prebiotic abundance (18–23) and their ability to enhance nonenzymatic copying at reduced concentrations. We found that 5 mM mononucleotides in combination with 1 mM trinucleotides or 0.5 mM tetranucleotides can also be nearly completely converted to either activated or bridged species (Figure 3C, Table S1). We compared these results with a control experiment where we observed the spontaneous formation of Ap*pCGCA from a mixture of pre-activated *pA and *pCGCA in the same primer extension buffer for 1 h (this is the optimal incubation period as shown in Figure S5). Despite starting with pure activated mono- and oligonucleotides, this control experiment produced less monomer-bridged-oligonucleotides than seen in the presence of MeNC activation chemistry (Figure 3Ciii-iv).

**Figure 3.**
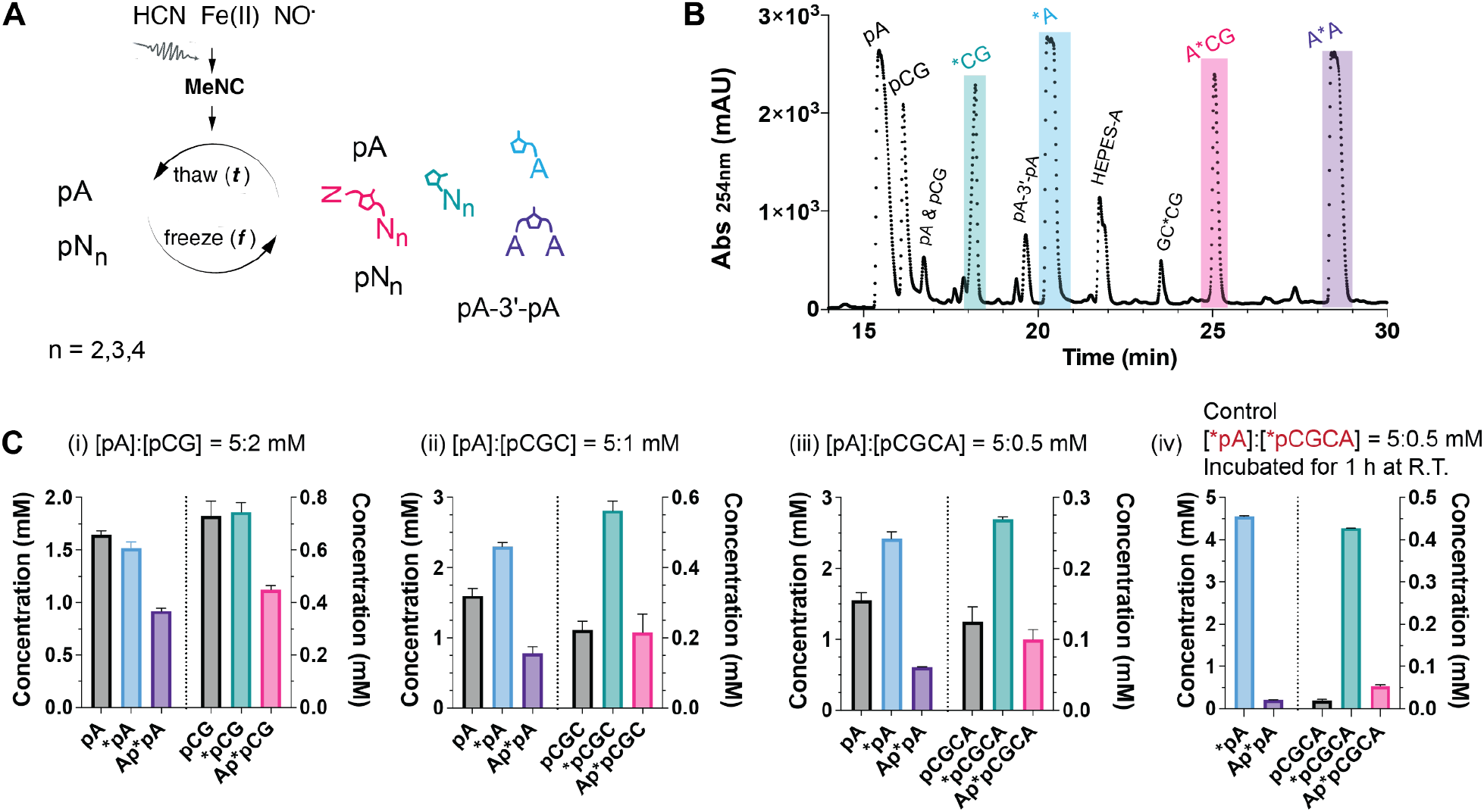
Ice-eutectic phase MeNC-mediated activation of oligonucleotides (*N_n_) and subsequent formation of monomer-bridged-oligonucleotides (N*N_n_). (A) A scheme illustrating the activation of mono- and oligonucleotides and the formation of bridged species (including N*N, N*N_n_, and N_n_*N_n_) in the ice-eutectic phase. (B) A representative HPLC trace of the purified end products from a reaction containing unactivated mono- and dinucleotides (5 mM pA and 2 mM pCG). (C) The concentration of each identified peak species from a reaction mixture of mono- and oligonucleotides of indicated length and concentration (i-iii) underwent eutectic phase activation, in contrast to the spontaneous formation of bridged species from a mixture of (iv) pure pre-activated mono- and tetranucleotides. Only a subset of the species is shown here for brevity (i-iii), but additional species can be found in Table S1. All activation reactions contained 30 mM MgCl_2_, 50 mM Na^+^-Hepes (pH 8) and stoichiometric 2AI ([2AI] = [pN]+[pN_n_]). 200 mM 2MBA and 50 mM MeNC were supplied to initiate the reaction, with subsequent periodic additions of MeNC in three aliquots of 50 mM at the beginning of each day of the freeze-thaw cycle. The corresponding elution profiles are shown in Figure S7. The control C (iv) was performed by incubating 5 mM *A, 0.5 mM *CGCA, 30 mM MgCl_2_, and 50 mM Na^+^-Hepes (pH 8) for 1 h at room temperature.

We then looked into whether a mixture of mononucleotides and short oligonucleotides of different lengths could be activated simultaneously and form bridged species. A mixture containing mono-, di-, and trinucleotides was activated efficiently with a substantial yield of bridged species (Figure S8). We monitored the reaction progress throughout the four days of activation and observed that the overall activation yields gradually increased while the monomer-bridged-oligonucleotide yield peaked on day two (Figure S8). It is encouraging that we were able to obtain mostly activated and bridged species from a heterogeneous mixture of mono- and oligonucleotides under a prebiotically plausible activation condition that is compatible with chemical RNA copying.

### *In-situ* activation of mononucleotides and oligonucleotides for nonenzymatic primer extension and ligation

Following the successful formation of monomer-bridged-oligonucleotides under the eutectic ice phase, we sought to apply this process to nonenzymatic RNA copying. We have previously demonstrated RNA copying driven by *in-situ* activation of mononucleotides, but we have not compared the process when both mono- and oligonucleotides are present, and when both primer extension and ligation can occur. In this study, we used a model system containing 5 mM pA and 0.5 mM pCGC to measure *in-situ* activation, primer extension, and ligation. To ensure an accurate comparison, we included control experiments using only the unactivated mononucleotide or only the unactivated oligonucleotide, thus preventing the formation of the monomer-bridged-oligonucleotide following activation. After the addition of MeNC and 2MBA, the samples were incubated under the eutectic phase (−15 to -13°C) and thawed every 24 hours for aliquot removal and the addition of fresh MeNC. After four days of eutectic activation, the samples were brought to room temperature for three more days of further extension. Additional MeNC and 2MBA were supplied in the last two days at room temperature to facilitate bridge formation, but efficient activation of nucleotides cannot occur at the low 2AI concentration we used (29, 30).

With the *in-situ* activation of pA and pCGC in the eutectic phase, we observed about a 70% yield of +1 primer extension within one day (Figure 4A). To determine whether the primer extension occurred during the eutectic ice phase or during the short thawing intervals, we quenched the 1-day reaction sample by adding EDTA before thawing, so that the Mg^2+^ would be chelated immediately upon thawing (Figure S9).

**Figure 4.**
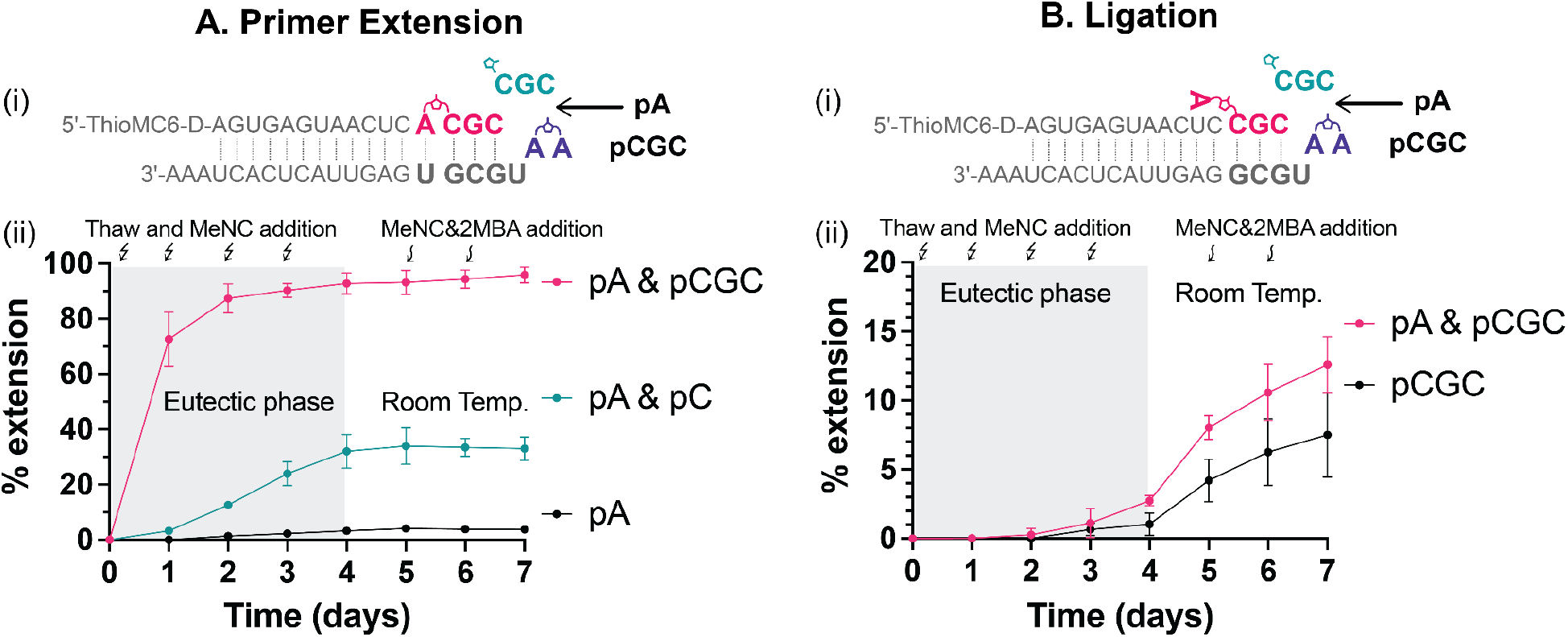
Ice-eutectic phase enables *in-situ* activation of mono- and oligonucleotides for enhanced nonenzymatic primer extension and ligation. (A) Primer extension with *in-situ* activated mono- and/or oligonucleotides. (i) Schematic representation of +1 primer extension with *in-situ* activated Ap*pCGC. (ii) Extension yield with: 5 mM pA (black), 5 mM pA & 5 mM pC (green), and 5 mM pA & 0.5 mM pCGC (pink). (B) Ligation with either *in-situ* activated oligonucleotides or a mixture of *in-situ* activated mono- and oligonucleotides. (i) Schematic representation of +3 ligation with *in-situ* activated Ap*pCGC. (ii) Ligation yield with: 0.5 mM pCGC (black) or 5 mM pA & 0.5 mM pCGC (pink). All reactions were initiated with 1 μM primer, 1.5 μM template, 5.5 mM 2AI, 50 mM Na^+^-Hepes (pH 8) and 30 mM MgCl_2_. 200 mM 2MBA was added at the beginning of the experiment, while 50 mM MeNC was freshly supplied every day. After four days of eutectic activation, the reactions were brought to ambient temperature for 24 h, then fresh 100 mM 2MBA and 100 mM MeNC were added every day for two more days.

We were surprised to find nearly identical extension yields with and without EDTA quenching prior to thawing. This result suggests that, despite the low temperature, the majority of the observed primer extension occurred in the ice eutectic phase. The considerable extent of primer extension reveals the efficient activation of mono- and oligonucleotides, as well as the high reactivity of the monomer-bridged-oligonucleotide even at low temperatures.

Since almost complete +1 primer extension was observed in one day with *in-situ* activated pA and pCGC, we further analyzed the extent of reaction throughout the first day by EDTA-quenching the partial frozen sample at different time points. We found that significant primer extension did not occur until after 6-8 h of eutectic phase activation (Figure S10). We interpret this result as indicating the time required for nucleotide activation and buildup of the monomer-bridged-oligonucleotide, before primer extension can happen.

In the case of a mixture of the mononucleotides pA and pC, and in the absence of oligonucleotides, the primer extension proceeded with sigmoidal behavior during the 4 days of eutectic activation, i.e. the reaction progressed slowly initially and accelerated at later times. This is likely because the concentration of Ap*pC peaked during the second day, consistent with our previous findings (29). However, we were surprised to see that primer extension almost completely halted during the subsequent incubation at ambient temperature (Figure 4Aii). We speculate that the substantial primer extension observed during the eutectic phase was due to the continuous generation of Ap*pC and the improved binding of Ap*pC to the template at low temperature. These benefits were lost once the reaction was brought to ambient temperature, leading to minimal subsequent primer extension. Similarly, primer extension when only pA was activated *in-situ* was very poor because the activation products, *pA and Ap*pA, could not bind well to the primer-template complex. These findings provide further support for the importance of utilizing oligonucleotides to catalyze template copying.

We also examined the ligation of *in-situ* activated pCGC to the same primer, using a different template. In contrast to the efficient +1 primer extension observed during the eutectic ice phase, the majority of this ligation did not take place until the room temperature incubation (Figure 4B). This difference is most likely explained by the low K_M_ and low k_obs max_ of the ligation reaction (Figure 1). Thus, the activated oligonucleotide can bind strongly to the template with multiple Watson-Crick base pairs, but the rate of the ligation reaction is intrinsically slow. As a result, ligation does not require the high effective concentration or better substrate binding attained under the ice eutectic phase. Instead, the low temperature of the eutectic phase suppresses the reaction. However, when the reaction is brought to room temperature, the ligation reaction can proceed slowly over 2-3 days. A greater extent of ligation was observed in conditions that favor the formation of monomer-bridged-oligonucleotides, likely due to the faster ligation rate with the monomer-bridged-oligonucleotide (Figure 1B).

Interestingly, in a typical nonenzymatic +1 primer extension reaction with *in-situ* activated pA and pCGC, the ligation product was also seen after the +A extension (Figures S11). This primer was first extended by +A using Ap*pCGC under the eutectic phase and then further extended by +CGC ligation at ambient temperature, likely using Ap*pCGC as well. These observations suggest that both primer extension and ligation may occur in an environment with freeze-thaw cycles followed by warm periods. For example, *in-situ* activation and primer extension could occur in the eutectic phase, followed by ligation after the temperature rises.

Since a prebiotically plausible reaction mixture would likely contain multiple short oligonucleotides, we also investigated *in-situ* activation and template copying with a mixture of pA, pACG, and pCGC. The pCGC can form Ap*pCGC to facilitate +1 primer extension, as previously demonstrated, while the pACG can be activated to *pACG and Ap*pACG, which are substrates for a competing ligation reaction. We found that the primer extension reaction was only slightly slower when the competing pACG was present (Figure S12). Almost no ligation products of pACG were observed, probably because most of the primer had already been rapidly extended by one nucleotide before +ACG ligation could happen. To simulate prebiotic conditions in which oligonucleotides were present at lower concentrations, we conducted competition experiments with decreasing concentrations of the two trinucleotides. We were surprised to find that template copying with only 50 μM of each trinucleotide behaved slightly better than the case with 500 μM of pACG and pCGC, possibly due to less inhibition from pACG, *pACG and Ap*pACG at lower concentrations and continuous generation of the highly reactive Ap*pCGC. Remarkably, substantial template copying was still observed even with trinucleotide concentrations as low as 5 μM each, indicating the strong catalytic effect of short oligonucleotides at minimal concentrations.

We have considered a prebiotic hot springs environment in which freeze-thaw cycles might be common. In such a scenario, mononucleotides and short oligonucleotides would be activated in the ice-eutectic phase, then periodically released by melting to flow to another site in which template copying could occur. To mimic this scenario, we first activated a mixture of mono- and oligonucleotides in the ice-eutectic phase as described above, but without the primer and the template. After one day of activation, the mixture was thawed and added to the primer-template complex. We were pleased to observe ∼80% +1 primer extension products in just 1 h (Figure 5). This nonenzymatic primer extension was performed at a reduced MgCl_2_ concentration of 30 mM (compared to 100 mM MgCl_2_ in Figures 1 and 2) to minimize hydrolysis, yet good template copying was still observed. Our findings suggest that *in-situ* activation chemistry and short oligonucleotides may have played an essential role in enhancing prebiotic nonenzymatic template copying.

**Figure 5.**
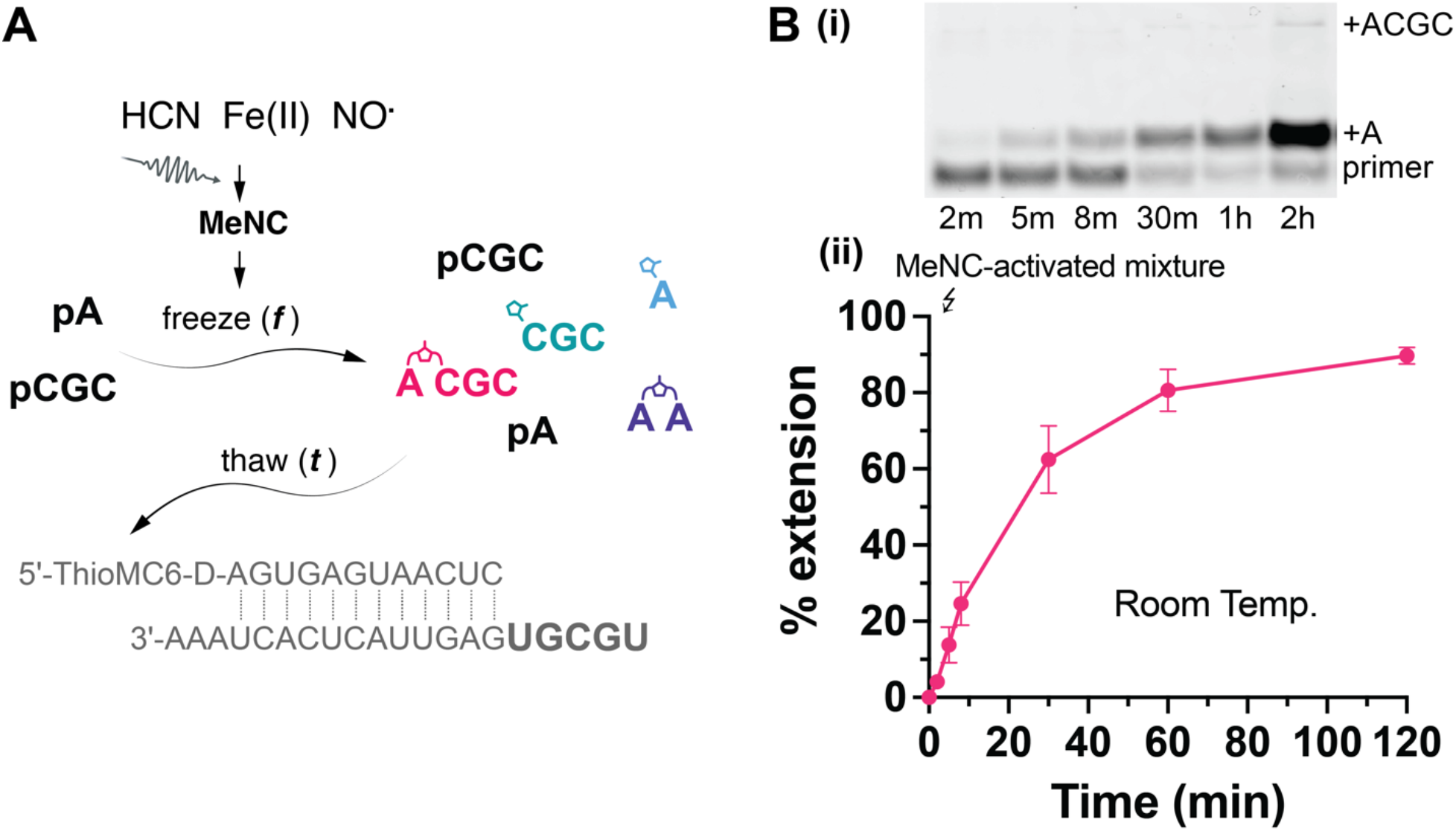
Nonenzymatic primer extension with pA and pCGC pre-activated in an ice-eutectic phase for 1 day. (A) A scheme illustrating the activation of pA and pCGC by MeNC-mediated chemistry in the ice-eutectic phase and the flow of materials to a new environment containing the primer-template complex upon thawing. (B) Primer extension with MeNC-activated mixture demonstrating ∼ 80% yield in 1 hour. (i) A representative PAGE gel analysis. (ii) Primer extension yield vs. time. The eutectic phase activation was performed for 1 day with 5 mM pA, 0.5 mM pCGC, 5.5 mM 2AI, 50 mM Na^+^-Hepes (pH 8), 30 mM MgCl_2_, 200 mM 2MBA, and 50 mM MeNC. After thawing, the mixture was added to the lyophilized primer-template complex to give a final reaction mixture containing 1 μM primer and 1.5 μM template.

## DISCUSSION

Any prebiotically plausible scenario for the nonenzymatic replication of RNA must include an efficient pathway for nucleotide activation, both initially and to counteract the inevitable hydrolysis of activated species. In addition, minimizing the complexity of the geochemical context is important, for example, by showing that multiple steps in a pathway can occur together in a single environment. We have now shown that freeze-thaw cycles can drive efficient activation of both mono- and oligonucleotides through the methyl isocyanide pathway. This efficient activation leads to the spontaneous synthesis of monomers that are imidazolium-bridged to short oligonucleotides, which in turn leads to efficient template-directed primer extension. Early work on nonenzymatic template copying required very high concentrations of activated monomers, typically in the range of 100 mM (9). In contrast, our work has shown that low mM concentrations of monomers together with micromolar concentrations of short oligonucleotides can lead to efficient copying by primer extension, which seems more likely in a prebiotic scenario. Below we discuss the implications of our results for the synthesis and catalytic roles of monomer-bridged-oligonucleotides in RNA replication, and we consider geochemical environments that could potentially host such RNA-catalyzed RNA replication processes.

Short oligonucleotides may have been generated on the primordial Earth via untemplated polymerization, e.g., on clay mineral surfaces or as a result of concentration in the ice eutectic phase of partially frozen water (18–20). Short oligonucleotides can also assemble by the template-directed oligomerization of activated monomers or bridged dinucleotides (21–23, 33). However, in the context of nonenzymatic copying, oligonucleotides are typically thought of as potential primers, templates, or ligators, and their potential roles as catalysts have not yet been fully explored. We have previously shown that monomer-bridged-oligonucleotide intermediates can significantly improve the rate of nonenzymatic primer extension (13). These substrates can spontaneously form in mixtures of activated mono- and oligonucleotides. We have now shown that isocyanide chemistry can also activate oligonucleotides of varying lengths, and thereby yield monomer-bridged-oligonucleotides under ice-eutectic conditions. Following the *in-situ* activation of monomers and oligonucleotides, we observed significant primer extension while in the ice-eutectic phase; we also observed considerable ligation but only following thawing. Furthermore, even at extremely low concentrations of short oligonucleotides (as low as 5 μM), nonenzymatic template copying driven by *in-situ* activation can be observed. Finally, we also demonstrated that short oligonucleotides can be activated in one environment and then transferred into a second environment where fast template copying can occur. Thus, short oligonucleotides may have played an important role in the catalysis of template copying on the primordial Earth. We are currently investigating the activation of complex mixtures of short oligonucleotides as a potential means of copying arbitrary RNA sequences.

We have recently proposed and provided the first experimental tests of a model for nonenzymatic RNA replication (33, 34). This virtual circular genome model invokes a collection of linear oligonucleotides of varying lengths that map onto both strands of a circular genomic sequence. In this model, all of the component oligonucleotides can play multiple roles, i.e., as primers, templates, and downstream helpers. Remarkably, activating all of the oligonucleotide components in this system significantly enhanced the observed rate of primer extension (34), consistent with a role for short, activated oligonucleotides in the catalysis of RNA copying chemistry. The combination of the prebiotically plausible activation chemistry presented in our current work with the virtual circular genome replication model has the potential to drive a nonenzymatic replication system through continuous *in-situ* activation.

It is interesting to consider whether the ice-eutectic mediated synthesis of bridged substrates could support RNA replication within protocells. Freeze-thaw cycles appear to be incompatible with vesicle integrity since ice crystals are thought to rupture membrane structures, resulting in the loss of compartmentalized content. We suggest that activation chemistry could happen in a nearby and partially frozen environment, where periodic thawing allows activated nucleotides to be released and flow into a separate environment in which protocells are replicating. Although monomer-bridged-oligonucleotide substrates are too large to readily diffuse across membranes and into protocells, we have recently described a phenomenon akin to spontaneous endocytosis in model protocells (35). Substrates internalized by endocytosis could then slowly diffuse into the protocell ‘cytoplasm’ on a longer time scale. An alternative possibility, which we are currently investigating, is that a modification of the isocyanide activation chemistry might obviate the need for freeze-thaw cycles, and allow for nucleotide and oligonucleotide activation to proceed within replicating protocells.

## SUPPLEMENTARY DATA

Supplementary data are available online.

## ACKNOWLEDGEMENTS

We thank Prof. Lijun Zhou and Dr. Saurja DasGupta for helpful discussions and comments on the manuscript.

## Reference

1. Joyce, G. F. (1989). RNA evolution and the origins of life. Nature, 338, 217–224.

2. Orgel, L.E. (2004) Prebiotic Chemistry and the Origin of the RNA World. Crit. Rev. Biochem. Mol. Biol., 39, 99–123.

3. Joyce, G. F., & Szostak, J. W. (2018). Protocells and RNA self-replication. Cold Spring Harb. Perspect. Biol., 10, a034801.

4. Wachowius, F., Attwater, J. and Holliger, P. (2017) Nucleic acids: function and potential for abiogenesis. Q. Rev. Biophys., 50, e4.

5. Becker, S., Feldmann, J., Wiedemann, S., Okamura, H., Schneider, C., Iwan, K., Crisp, A., Rossa, M., Amatov, T. and Carell, T. (2019). Unified prebiotically plausible synthesis of pyrimidine and purine RNA ribonucleotides. Science, 366, 76–82.

6. Patel, B.H., Percivalle, C., Ritson, D.J., Duffy, C.D. and Sutherland, J.D. (2015) Common origins of RNA, protein and lipid precursors in a cyanosulfidic protometabolism. Nat. Chem. 2015 7:4, 7, 301– 307.

7. Weimann, B.J., Lohrmann, R., Orgel, L.E., Schneider-Bernloehr, H. and Sulston, J.E. (1968) Template-Directed Synthesis with Adenosine-5′-phosphorimidazolide. Science (1979), 161, 387.

8. Sulston, J., Lohrmann, R., Orgel, L.E. and Miles, H.T. (1968) Nonenzymatic synthesis of oligoadenylates on a polyuridylic acid template. Proc. Natl. Acad. Sci., 59, 726–733.

9. Rembold, H., & Orgel, L. E. (1994). Single-strand regions of Poly (G) act as templates for oligo (C) synthesis. J. Mol. Evol., 38, 205–210.

10. Walton, T. and Szostak, J.W. (2016) A Highly Reactive Imidazolium-Bridged Dinucleotide Intermediate in Nonenzymatic RNA Primer Extension. J. Am. Chem. Soc., 138, 11996–12002.

11. Walton, T. and Szostak, J.W. (2017) A Kinetic Model of Nonenzymatic RNA Polymerization by Cytidine-5′-phosphoro-2-aminoimidazolide. Biochemistry, 56, 5739–5747.

12. Walton, T., Zhang, W., Li, L., Tam, C.P. and Szostak, J.W. (2019) The Mechanism of Nonenzymatic Template Copying with Imidazole-Activated Nucleotides. Angew. Chem. Int. Ed., 58, 10812–10819.

13. Ding, D., Zhou, L., Giurgiu, C. and Szostak, J.W. (2022) Kinetic explanations for the sequence biases observed in the nonenzymatic copying of RNA templates. Nucleic Acids Res., 50, 35–45.

14. Zhang, W., Tam, C.P., Walton, T., Fahrenbach, A.C., Birrane, G. and Szostak, J.W. (2017) Insight into the mechanism of nonenzymatic RNA primer extension from the structure of an RNA-GpppG complex. Proc. Natl. Acad. Sci., 114, 7659–7664.

15. Zhang, W., Walton, T., Li, L. and Szostak, J.W. (2018) Crystallographic observation of nonenzymatic RNA primer extension. Elife, 7, e36422.

16. Duzdevich, D., Carr, C.E., Ding, D., Zhang, S.J., Walton, T.S. and Szostak, J.W. (2021) Competition between bridged dinucleotides and activated mononucleotides determines the error frequency of nonenzymatic RNA primer extension. Nucleic Acids Res, 49, 3681–3691.

17. Prywes, N., Blain, J.C., del Frate, F. and Szostak, J.W. (2016) Nonenzymatic copying of RNA templates containing all four letters is catalyzed by activated oligonucleotides. Elife, 5.

18. Ferris, J.P. and Ertem, G. (1993) Montmorillonite Catalysis of RNA Oligomer Formation in Aqueous Solution. A Model for the Prebiotic Formation of RNA. J. Am. Chem. Soc., 115, 12270–12275.

19. Kanavarioti, A., Monnard, P.A. and Deamer, D.W. (2001) Eutectic phases in ice facilitate nonenzymatic nucleic acid synthesis. Astrobiology, 1, 271–281.

20. Monnard, P.A., Kanavarioti, A. and Deamer, D.W. (2003) Eutectic Phase Polymerization of Activated Ribonucleotide Mixtures Yields Quasi-Equimolar Incorporation of Purine and Pyrimidine Nucleobases. J. Am. Chem. Soc., 125, 13734–13740.

21. Lohrmann, R., Bridson, P.K. and Orgel, L.E. (1980) Efficient Metal-Ion Catalyzed Template-Directed Oligonucleotide Synthesis. Science (1979), 208, 1464–1465.

22. Inoue, T. and Orgel, L.E. (1982) Oligomerization of (guanosine 5′-phosphor)-2-methylimidazolide on poly(C). An RNA polymerase model. J. Mol. Biol., 162, 201–217.

23. Trinks, H., Schröder, W. and Biebricher, C.K. (2005) Ice and the origin of life. Orig. Life. Evol. Biosph., 35, 429–445.

24. Rohatgi, R., Bartel, D. P., & Szostak, J. W. (1996). Kinetic and mechanistic analysis of nonenzymatic, template-directed oligoribonucleotide ligation. J. Am. Chem. Soc., 118, 3332–3339.

25. Zhou, L., O’Flaherty, D.K. and Szostak, J.W. (2020) Template-Directed Copying of RNA by Non-enzymatic Ligation. Angew. Chem., 132, 15812–15817.

26. Zhou, L., O’Flaherty, D.K. and Szostak, J.W. (2020) Assembly of a Ribozyme Ligase from Short Oligomers by Nonenzymatic Ligation. J. Am. Chem. Soc., 142, 15961–15965.

27. Wachowius, F. and Holliger, P. (2019) Non-Enzymatic Assembly of a Minimized RNA Polymerase Ribozyme. ChemSystemsChem, 1, 12–15.

28. Mariani, A., Russell, D.A., Javelle, T. and Sutherland, J.D. (2018) A Light-Releasable Potentially Prebiotic Nucleotide Activating Agent. J. Am. Chem. Soc., 140, 8657–8661.

29. Zhang, S.J., Duzdevich, D., Ding, D. and Szostak, J.W. (2022) Freeze-thaw cycles enable a prebiotically plausible and continuous pathway from nucleotide activation to nonenzymatic RNA copying. Proc. Natl. Acad. Sci., 119, e2116429119.

30. Zhang, S.J., Duzdevich, D. and Szostak, J.W. (2020) Potentially prebiotic activation chemistry compatible with nonenzymatic RNA copying. J. Am. Chem. Soc., 142, 14810–14813.

31. Zhang, W., Tam, C.P., Zhou, L., Oh, S.S., Wang, J. and Szostak, J.W. (2018) Structural Rationale for the Enhanced Catalysis of Nonenzymatic RNA Primer Extension by a Downstream Oligonucleotide. J. Am. Chem. Soc., 140, 2829–2840.

32. Giurgiu, C., Fang, Z., Aitken, H. R., Kim, S. C., Pazienza, L., Mittal, S., & Szostak, J. W. (2021). Structure–Activity Relationships in Nonenzymatic Template-Directed RNA Synthesis. Angew. Chem. Int. Ed. Engl., 60, 22925–22932.

33. Zhou, L., Ding, D. and Szostak, J.W. (2021) The virtual circular genome model for primordial RNA replication. RNA, 27, 1–11.

34. Ding, D., Zhou, L., Mittal, S. and Szostak, J.W. (2023) Experimental Tests of the Virtual Circular Genome Model for Non-enzymatic RNA Replication. J. Am. Chem. Soc., 145, 7504–7515

35. Stephanie, J. Z., Lauren, A. L., Palapuravan, A., Yamuna, K., Thomas, G. F., Jack, W. S., and Anna, W. (2023). Passive endocytosis in model protocells. bioRxiv, 2023.2001.2007.522792. doi:10.1101/2023.01.07.522792

