## Supplementary Information for "Enhanced nonenzymatic RNA copying with *in-situ* activation of short oligonucleotides"

**Table of Contents**

1. Supplementary Figures S1 to S12 S3-14

2. Supplementary Table S1 S15

3. Supplementary References S15

**Supplementary Figures:**

**Figure S1**

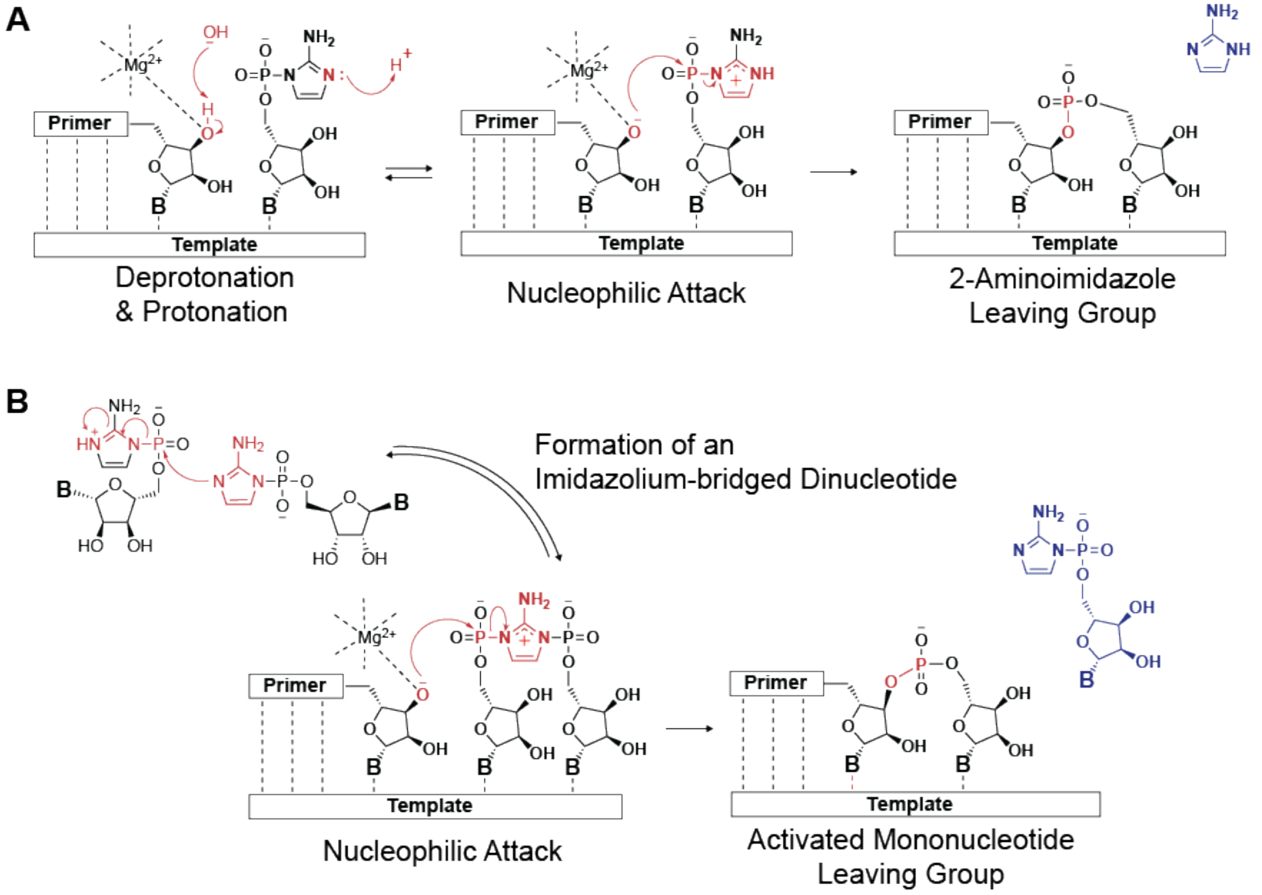


**Figure S1. Schematic representation of the presumed mechanism of primer extension by one nucleotide.** (A) Nonenzymatic primer extension using a 2-aminoimidazole activated mononucleotide. (B) Nonenzymatic primer extension using a 2-aminoimidazolium bridged dinucleotide.

**Figure S2**

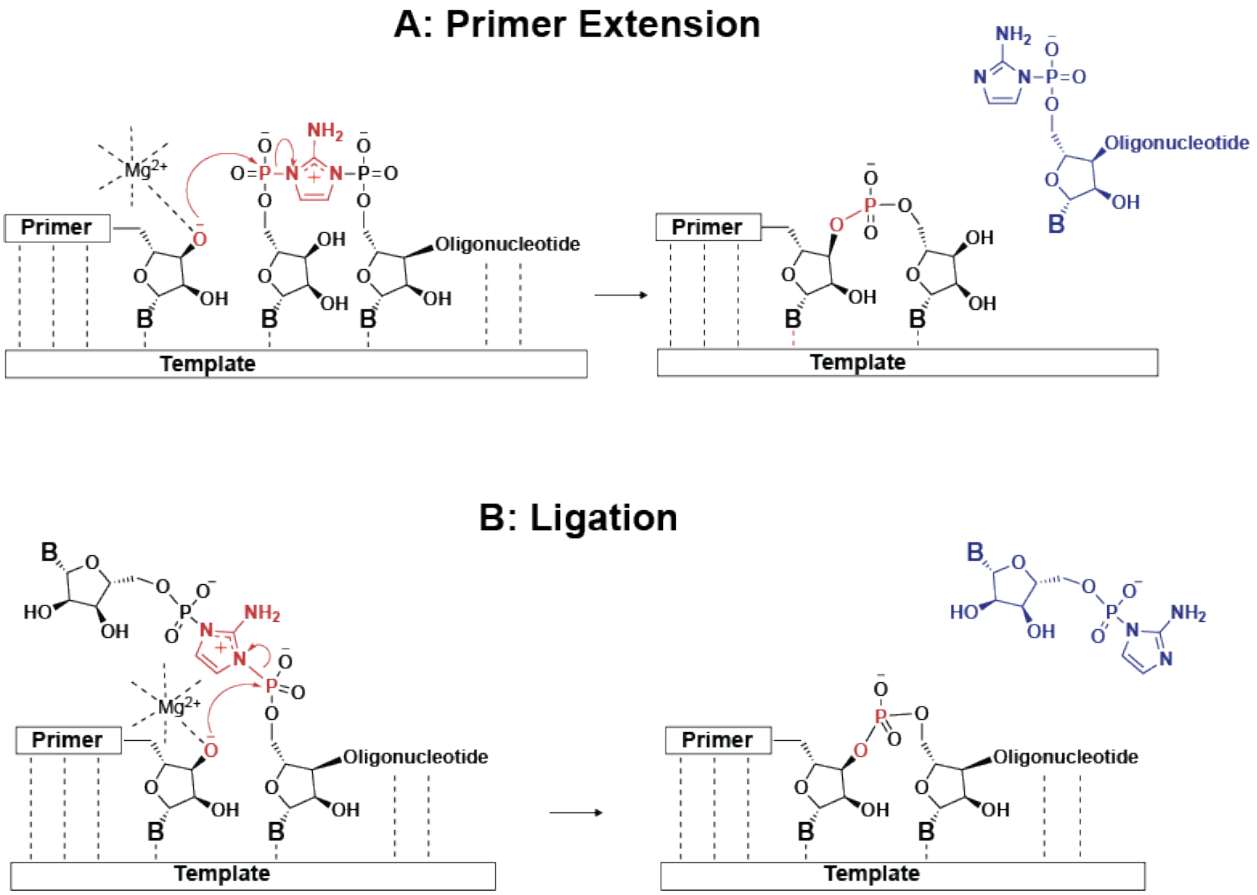


**Figure S2. Scheme for nonenzymatic template copying with a monomer-bridged-oligonucleotide.** (A) Primer extension of one nucleotide and displacement of a 2AI-oligonucleotide as the leaving group. (B) Ligation of an oligonucleotide and displacement of a 2AI-mononucleotide as the leaving group.

**Figure S3**

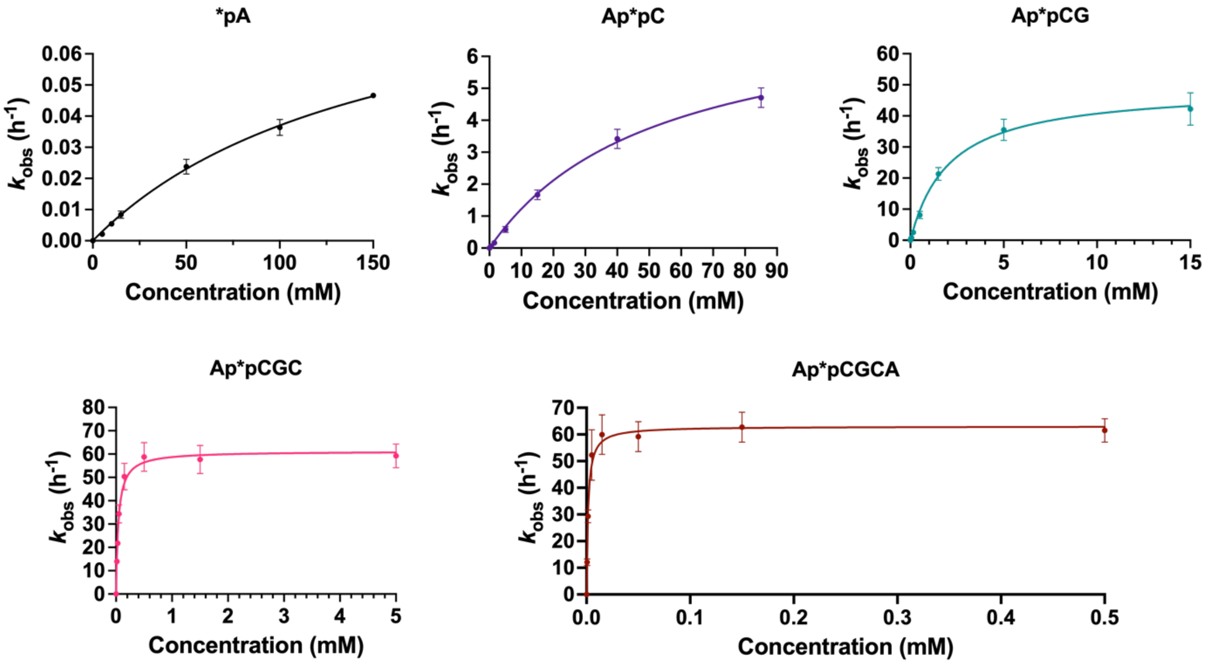


**Figure S3. Michaelis-Menten curves for primer extension with *pA, Ap*pC, Ap*pCG, Ap*pCGC, and Ap*pCGCA.** Primer extension experiments were performed with 1.5 µM primer, 2.5 µM template, 100 mM $MgCl_2$ and 200 mM Tris-HCl (pH 8.0). Modified from Fig. S5 in ref. 1 with permission under a Creative Commons Attribution 4.0 International License. Copyright 2022 Ding et al.; Published by Oxford University Press on behalf of Nucleic Acids Research.

### Figure S4

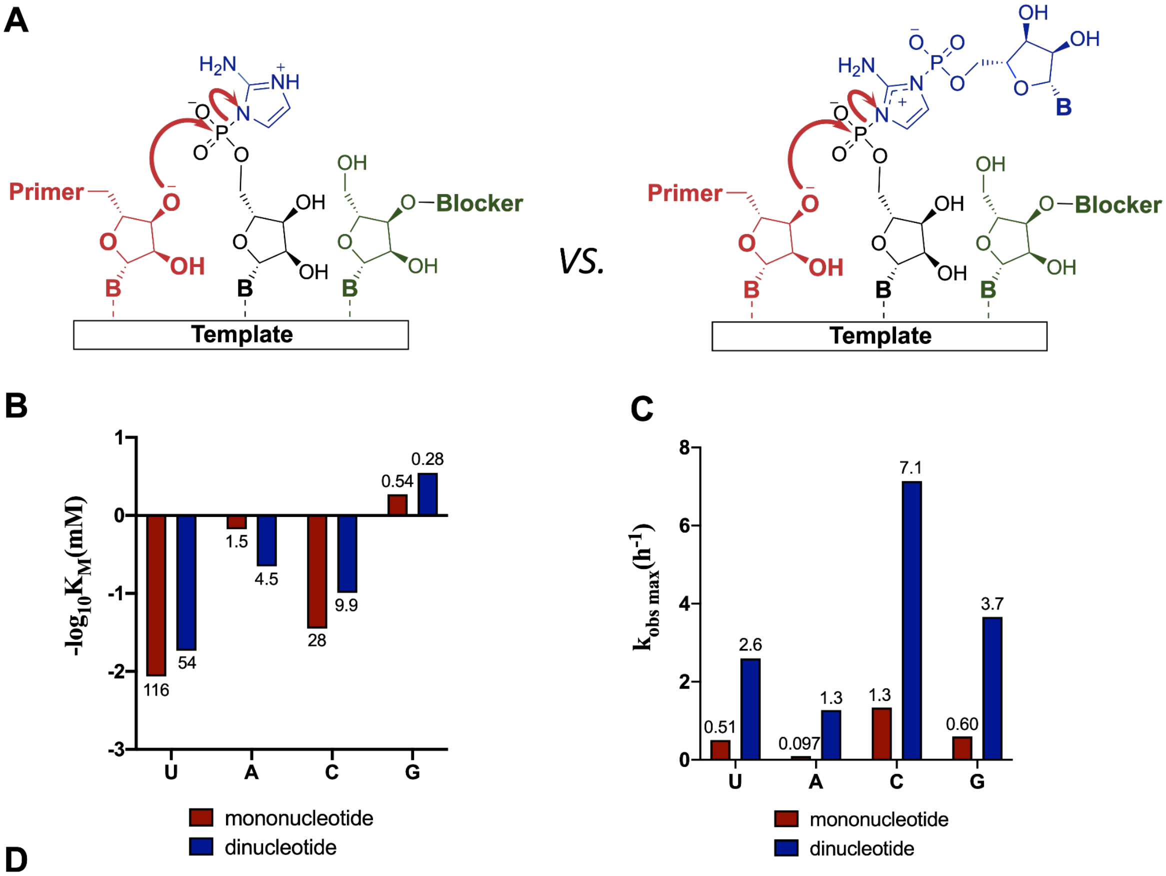


D

| Activated substrate | Primer 5'FAM->3' | Template 5'OH->3' | Blocker 5'OH->3' |
|---|---|---|---|
| *pU/Up*pU | /FAM/AGUGAGUAACGG | AUAUCUGACAUCC A CCGUUACUCACU | GG AUG UCA GAU AU |
| *pA/Ap*pA | /FAM/AGUGAGUAACGG | AUAUCUGACAUCC U CCGUUACUCACU | GG AUG UCA GAU AU |
| *pC/Cp*pC | /FAM/AGUGAGUAACGG | AUAUCUGACAUCC G CCGUUACUCACU | GG AUG UCA GAU AU |
| *pG/Gp*pG | /FAM/AGUGAGUAACGG | AUAUCUGACAUCC C CCGUUACUCACU | GG AUG UCA GAU AU |

**Figure S4. Comparison between 2-aminoimidazole activated mononucleotide and 2-aminoimidazolium bridged dinucleotide for primer extension by one nucleotide with single nucleotide binding site.** (A) Schematic representation for the primer-template-blocker system with 1-nt binding site and the mechanism for +1 extension either through activated mononucleotide or bridged dinucleotide. (B) Comparison of the binding affinity with either *pN or Np*pN. (C) Comparison of the maximum reaction rate at saturating binding with either *pN or Np*pN. (D) Sequences for the primer, template, and blocker used for the assay. The underlined letters in the template sequences indicate the nucleotide at each open binding site. All reactions were performed with 1.5 µM primer, 2.5 µM template, 3.5 µM blocker, 100 mM $MgCl_2$ and 200 mM Tris-HCl (pH 8.0). The downstream blocker nucleotide is not activated and only serves to block downstream binding sites. $K_M$ and $k_{obs\ max}$ were measured by fitting primer extension rates with respect to concentrations to the Michalis-Menten equation. The exact values of $K_M$ (mM) and $k_{obs\ max}$ ($h^{-1}$) are indicated on the top or at the bottom of each bar.

**Figure S5**

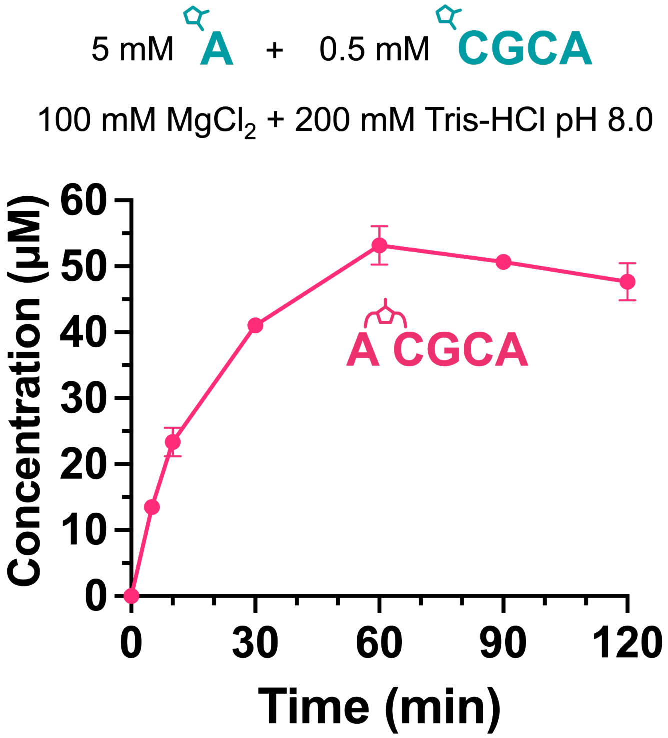


**Figure S5. HPLC quantification of Ap*pCGCA spontaneously formed in solution.** 5 mM *pA and 0.5 mM *pCGCA were incubated in 100 mM $MgCl_2$ and 200 mM Tris-HCl pH 8.0. The concentration of Ap*pCGCA peaked at 1 h, after which the Ap*pCGCA was slowly hydrolyzed.

**Figure S6**

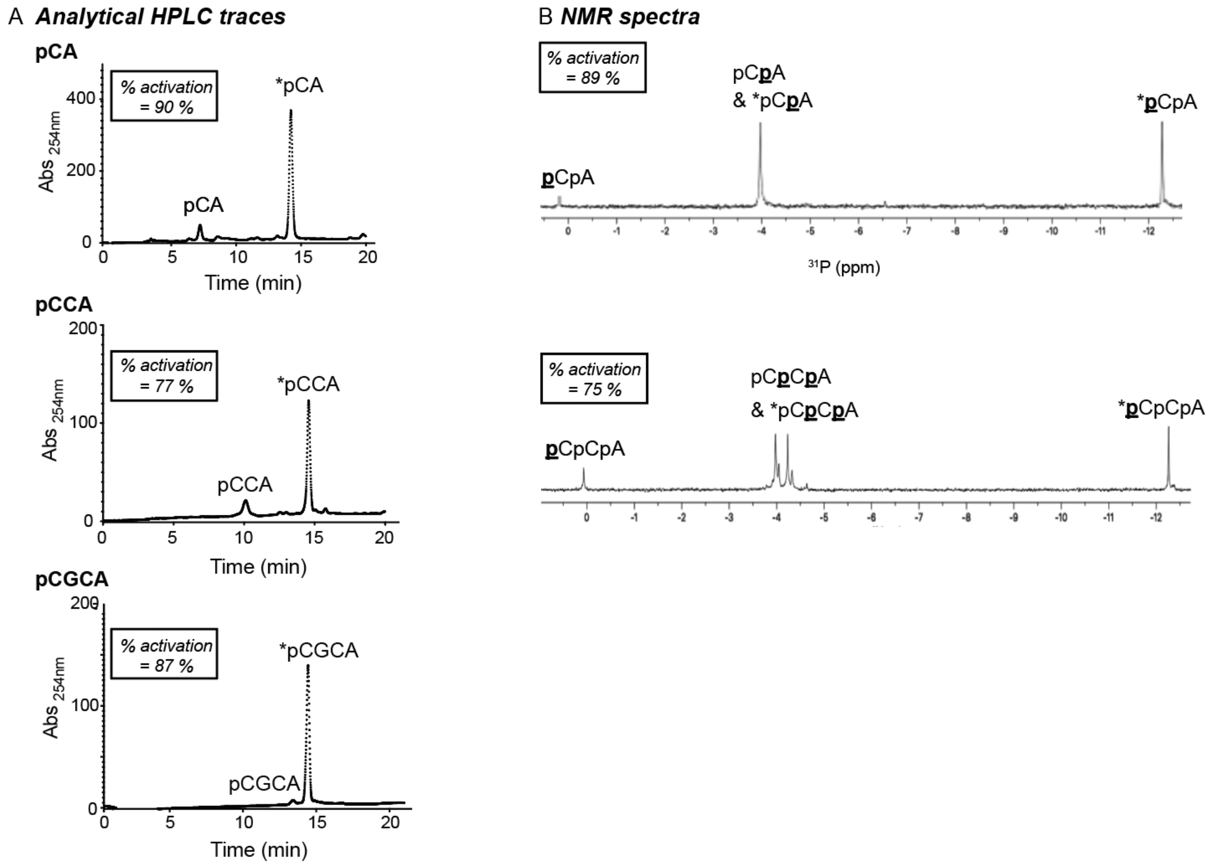


**Figure S6. Quantification of the isocyanide-mediated phosphate activation of varying oligomers with 2AI to demonstrate the consistency between the NMR and HPLC measurements.** (A) Representative HPLC traces at t = 6 h showing 5′-monophosphorylated oligonucleotides and 2AI-activated oligonucleotides, e.g., elution peaks at 7.4 and 14.3 min correspond to 5′-monophosphorylated dinucleotide (pCA) and 2AI-activated dinucleotide (*pCA), respectively. (B) $^{31}P$ NMR spectra of 2AI-activated oligonucleotides formed by isocyanide chemistry at 6 h. Corresponding phosphorus nucleus/nuclei are highlighted in bold and underlined. Quantification of the 2AI-activated activation yields by NMR and HPLC spectroscopy has demonstrated the consistency between the two different measuring methods. Reaction conditions: 400 mM MeNC, 400 mM 2MBA, and 200 mM 2AI were added to the solution of 5 mM oligonucleotides in 200 mM $Na^+$-Hepes at pH 8 with 30 mM $MgCl_2$. The reaction was allowed to sit for 6 hours, which was the optimal incubation time as determined previously (2), at ambient temperature. MeNC was removed before analysis by HPLC by Sep-Pak® C18 Cartridge chromatography. Samples were adjusted to 10 % (v/v) $D_2O$ prior to NMR spectroscopy.

**Figure S7.**

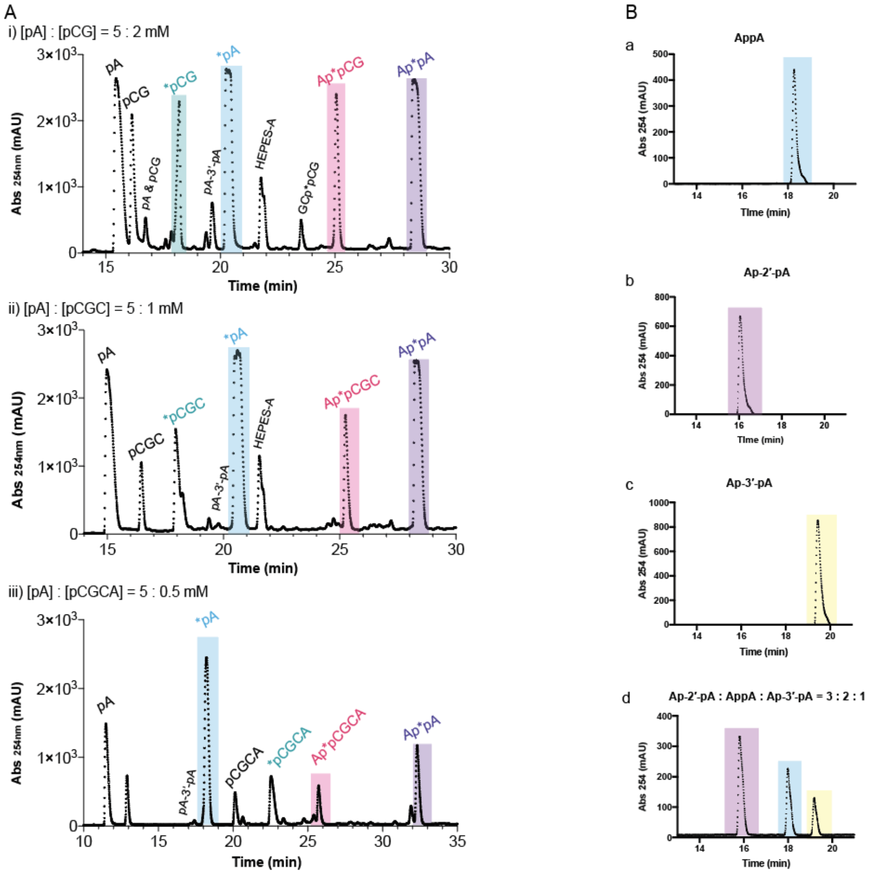

**Figure S7. Products of activation reactions after freeze-thaw cycles.** (A) Representative HPLC traces of the products of activation chemistry after removal of isocyanide from i) unactivated mono- and di-nucleotides (5 mM pA and 2 mM pCG), ii) mono- and trinucleotides (5 mM pA and 1 mM pCGC), and iii) mono- and tetra-nucleotides (5 mM pA and 0.5 mM pCGCA), following eutectic phase MeNC activation as shown in Figure 3. (B) HPLC traces of authentic synthetic (a) 5′-5′-pyrophosphate di-adenosine (AppA), (b) di-adenosine monophosphate with 2′-5′ phosphodiester linkages (pA-2′-pA), (c) di-adenosine monophosphate with 3′-5′ phosphodiester linkages (pA-3′-pA), and (d) a mixture of AppA, pA-2′-pA, and pA-3′-pA in a ratio of 2:3:1. The elution positions of these compounds were used to help identify peaks in part (A).

HPLC conditions: All samples were separated by HPLC as described in detail in the Methods (2 % to 13 % ACN in TEAB buffer at pH 8 over 35 minutes), except that the products from the mixture of mono- and tetra-nucleotides (A iii) were eluted between 2 % and 11 % ACN in TEAB buffer at pH 8 over 35 minutes.

Reaction conditions: activation reactions in (A) contained stoichiometric 2AI ([2AI] = [pN]+[pN$_n$]), 200 mM 2MBA, 50 mM MeNC, 30 mM $MgCl_2$, 50 mM $Na^+$-Hepes at pH 8, and subsequent periodic addition of MeNC in three aliquots of 50 mM at the beginning of each freeze-thaw cycle.

**Figure S8**

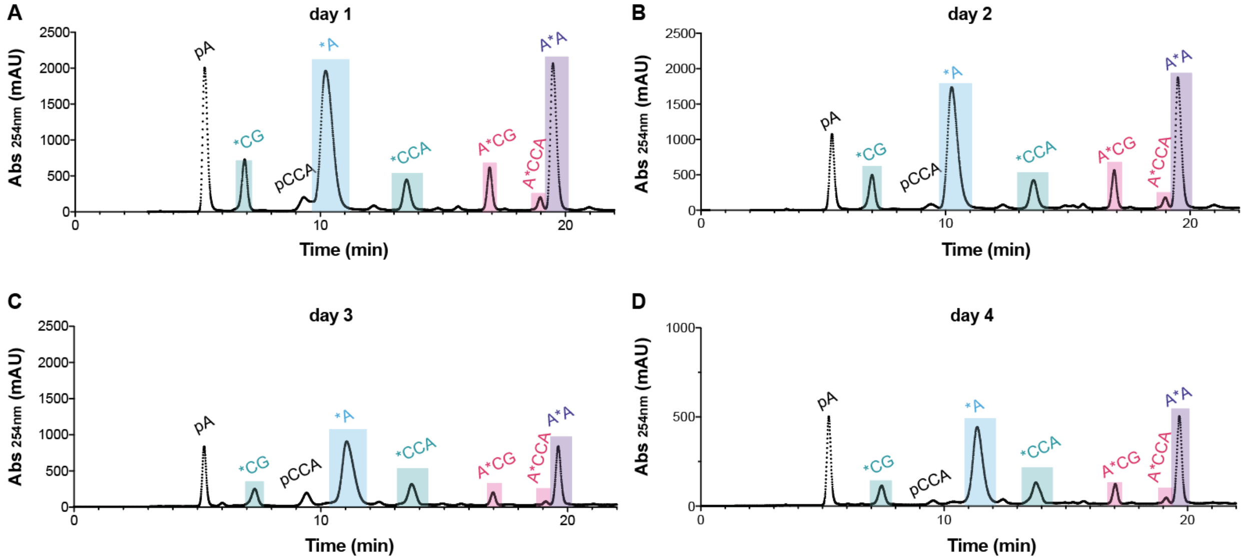


**Figure S8. HPLC analysis of the products of an ice eutectic phase activation reaction of a mixture of unactivated mono-, di- and trinucleotides (5 mM pA, 1 mM pCG, and 0.5 mM pCCA).** Activation reactions contained 6.5 mM 2AI ([2AI] = [pN]+[$pN_n$]), 200 mM 2MBA, 50 mM MeNC, 30 mM $MgCl_2$, 50 mM $Na^+$-Hepes pH 8, with subsequent periodic addition of MeNC in three aliquots of 50 mM at the beginning of each daily freeze-thaw cycle. Post-reaction cleanup using Sep-Pak® C18 Cartridges was performed to remove MeNC prior to HPLC analysis. The reaction products were eluted between 2 % and 13 % ACN in TEAB buffer at pH 8 over 25 minutes.

**Figure S9**

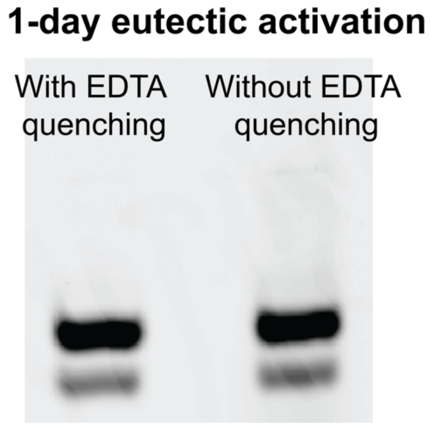


**Figure S9. Comparison between primer extension reactions with and without EDTA quenching in the ice eutectic phase.** (Left) EDTA quenching of 1-day primer extension products with *in-situ* eutectic activation of pA and pCGC. To ensure any thawed component is thoroughly quenched, the reaction at frozen eutectic phase was treated directly with excess EDTA and extensive vortexing. (Right) A parallel reaction under the identical condition, except without the EDTA quenching before thawing. Both reactions were performed with 5 mM pA, 0.5 mM pCGC, 5.5 mM 2AI, 50 mM $Na^+$-Hepes at pH = 8.0 and 30 mM $MgCl_2$. The mixture was brought to 200 mM 2MBA and 50 mM MeNC before being kept at -15° C for 24 h.

**Figure S10**

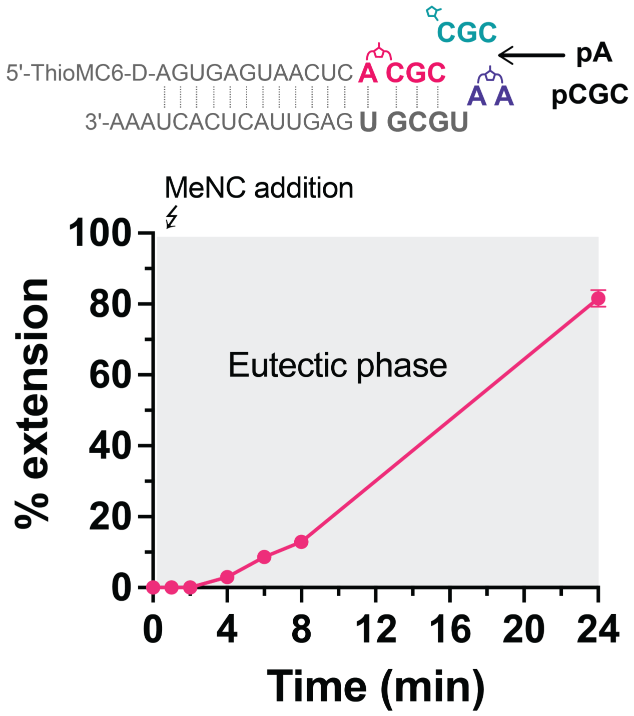


**Figure S10. Time course of nonenzymatic primer extension with *in-situ* activation of pA and pCGC under ice eutectic conditions.** Each sample was prepared separately and quenched with EDTA before thawing. Each reaction was performed with 5 mM pA, 0.5 mM pCGC, 5.5 mM 2AI, 50 mM $Na^+$-Hepes at pH = 8.0 and 30 mM $MgCl_2$. The mixture was brought to 200 mM 2MBA and 50 mM MeNC before being incubated at -15° C.

**Figure S11**

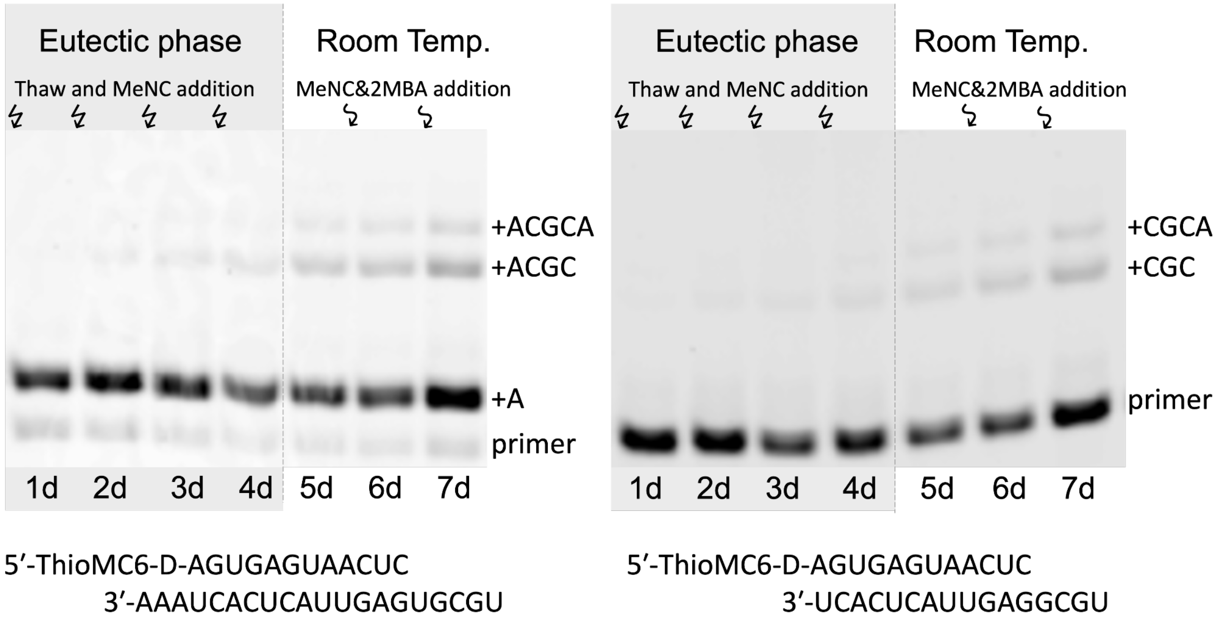


**Figure S11. PAGE gel analysis of primer-extension reactions with *in-situ* activation of 5 mM pA and 0.5 mM pCGC.** The primer and template sequences are indicated at the bottom. The identities of the primer-extension products are labeled at the right side of the gel. (Left) Primer extension as shown in Figure 4A with pA and pCGC. (Right) Ligation reaction as shown in Figure 4B with pA and pCGC. Both reactions were performed with 5.5 mM 2AI, 50 mM $Na^+$-Hepes (pH = 8.0) and 30 mM $MgCl_2$. 200 mM 2MBA was added at the beginning of the experiment while 50 mM MeNC was added at the beginning of every cycle of ice eutectic phase activation. After four cycles of eutectic activation, the reactions were brought to ambient temperature for 24 h, then fresh 100 mM 2MBA and 100 mM MeNC were added every day for two more days.

**Figure S12**

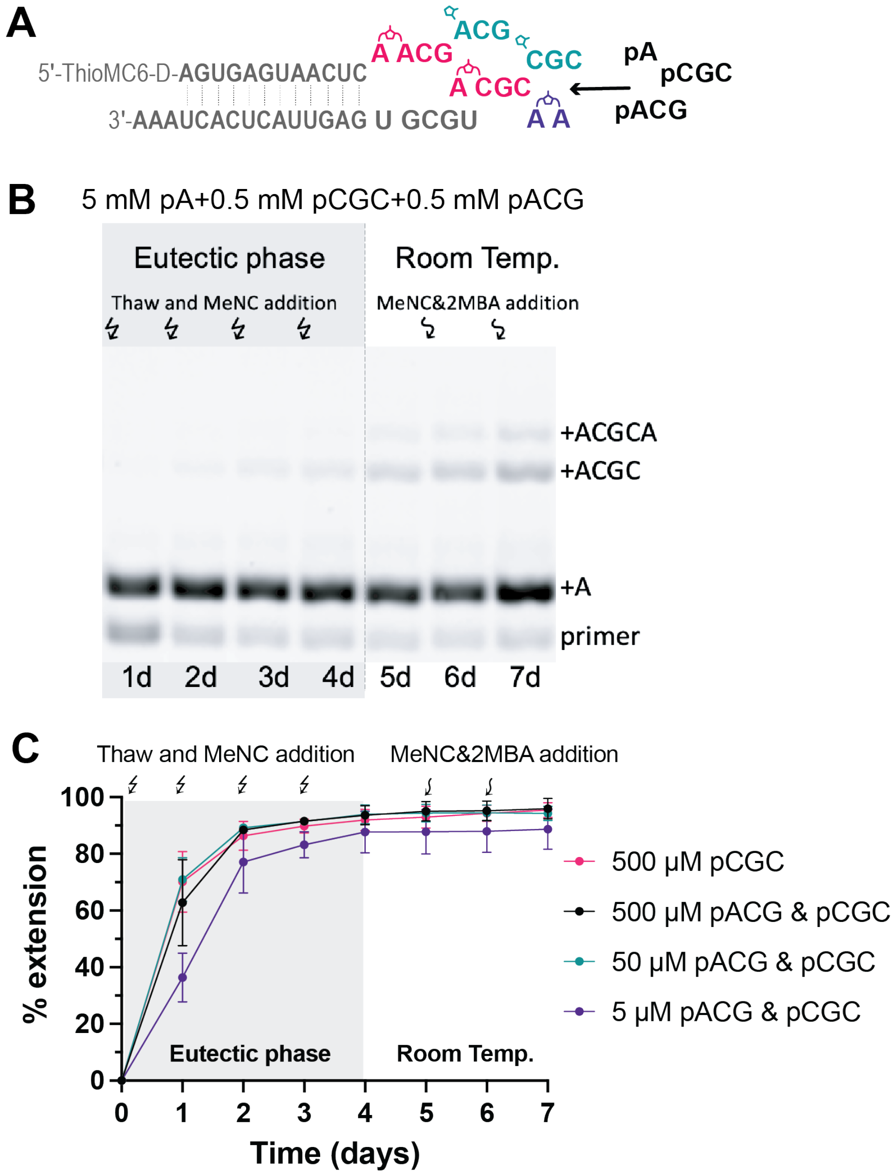


**Figure S12. Primer extension with 5 mM pA and oligonucleotide mixtures (pACG & pCGC) at decreasing concentrations.** (A) Schematic representation (B) A representative PAGE gel analysis of extension products with *in-situ* activation of 5 mM pA, 0.5 mM pCGC, and 0.5 mM pACG. (C) Primer extension yield with *in-situ* activation. All reactions were performed with 50 mM $Na^+$-Hepes (pH 8) and 30 mM $MgCl_2$. 200 mM 2MBA was added at the beginning of the experiment, while 50 mM MeNC was freshly supplied every day. After four days of eutectic activation, the reactions were brought to ambient temperature for 24 h, then fresh 100 mM 2MBA and 100 mM MeNC were added every day for two more days.

### Supplementary Table

**Table S1. The concentration[a] of each end product (mM) measured by HPLC[b] from Figure 3**

| [pA] : [pCG] = 5 : 2 mM | | [pA] : [pCGC] = 5 : 1 mM | | [pA] : [pCGCA] = 5 : 0.5 mM | |
|---|---|---|---|---|---|
| pA | 1.64± 0.03 | pA | 1.6±0.1 | pA | 1.6±0.1 |
| pCG | 0.73± 0.05 | pCGC | 0.22±0.03 | pCGCA | 0.13±0.02 |
| *pCG | 0.71 ± 0.02 | *pCGC | 0.56±0.03 | *pCGCA | 0.270±0.003 |
| pA-3′-pA[c] | 0.12 ± 0.02 | pA-3′-pA[c] | 0.034±0.003 | pA-3′-pA[c] | 0.14±0.11 |
| *A | 1.51 ± 0.06 | *pA | 2.30±0.06 | *pA | 2.43±0.09 |
| Hepes-A | 0.45 ± 0.03 | Hepes-A | 0.12±0.03 | Hepes-A | 0.16±0.09 |
| GCp*pCG | 0.07 ± 0.01 | Ap*pCGC | 0.22±0.05 | Ap*pCGCA | 0.10±0.01 |
| Ap*pCG | 0.45 ± 0.01 | Ap*pA | 0.78±0.09 | Ap*pA | 0.603±0.006 |
| Ap*pA | 0.92 ± 0.02 | | | | |

[a]The concentrations were determined from the normalized integrations of the elution peaks, where normalization is based on the UV absorbance of respective species. The concentration of each species was determined based on the concentrations of initial mononucleotides and oligonucleotides, and the ratio of normalized integrations between different species. [b]All reactions were carried out using 5 mM mononucleotides and the indicated amount and length of short oligonucleotides with stoichiometric 2AI ([2AI] = [pN]+[$(pN)_n$]), 200 mM 2MBA, 50 mM MeNC, 30 mM $MgCl_2$, 50 mM $Na^+$-Hepes pH 8, and subsequent periodic addition of MeNC in three aliquots of 50 mM. After the last addition of MeNC, the solution was left under eutectic ice phase conditions for 24 h. The end products were then separated from MeNC-mediated activation reagents and analyzed on HPLC. All peak fractions were flash frozen and lyophilized before confirming its identity by (LC-MS). [c] The identity of the $pA_2$ were determined by HPLC analysis of authentic sample AppA, pA-2′-pA, and pA-3′-pA (Figure S7).

### Supplementary References


1. Ding, D., Zhou, L., Giurgiu, C. and Szostak, J.W. (2022) Kinetic explanations for the sequence biases observed in the nonenzymatic copying of RNA templates. *Nucleic Acids Res.*, **50**, 35–45.

2. Zhang, S.J., Duzdevich, D. and Szostak, J.W. (2020) Potentially prebiotic activation chemistry compatible with nonenzymatic RNA copying. *J. Am. Chem. Soc.*, **142**, 14810–14813.
